# Uncovering Functional Gene Regulatory Networks in Bulk and Single-Cell Data through Robust Transcription Factor Activity Estimation and Model-Guided Experimental Validation

**DOI:** 10.1101/2025.06.09.658650

**Authors:** Alireza Fotuhi Siahpirani, Sunnie Grace McCalla, Saptarshi Pyne, Caleb Dillingham, Rupa Sridharan, Sushmita Roy

## Abstract

Reconstructing genome-scale gene regulatory networks (GRNs) remains a difficult problem in systems biology, and many experimental and computational methods have been developed to address this problem. Recent computational methods have aimed to more accurately model GRNs by estimating the hidden Transcription Factor Activity (TFA) from prior knowledge of TF target regulatory connections, encoded as an input directed graph, to relax the assumption that mRNA level of the regulator correlates with the protein activity of the regulator. However, the noise in the prior knowledge can adversely affect the estimated TFA levels and the quality of the downstream inferred GRNs. Here, we present a new approach, MERLIN+P+TFA, that uses prior knowledge-guided sparsity regularization to robustly and accurately estimate TFA and downstream GRNs. We apply our method to simulated and real expression data in yeast and mammalian systems and show improved quality of inferred GRNs for both bulk and single-cell datasets. Regularized TFA offers benefits to a variety of other GRN inference algorithms, including those that have traditionally been used with expression alone, in both bulk and scRNA-seq settings. We used the inferred GRN to prioritize key regulators for the mouse Embryonic Stem Cell (mESC) state and validate 58 regulators experimentally. We identified both known and novel regulators of the mESC state and further validate the targets of 4 known and novel regulators. Our validation experiments suggest that computationally inferred networks can capture functional targets of TFs with higher precision than estimated in current benchmarks, however, it is important to generate context-specific gold standards.

## Introduction

Transcriptional gene regulatory networks determine the context-specific expression levels of genes by specifying which regulatory proteins such as transcription factors and signaling proteins, regulate the expression of a gene. Expression-based network inference is a powerful way to reconstruct an initial genome-scale gene regulatory network. Given genome-wide mRNA profiles, these methods predict the regulators of a gene based on the ability of a regulator to explain the variation in the mRNA level of a target gene. Current methods for network inference can be grouped into those that rely on expression alone [1] and those that combine gene expression with other data sources, such as sequence-specific motif information, ChIP-seq datasets as prior knowledge to constrain the network structure [2–8]. A major limitation of these approaches is that mRNA levels of regulators are assumed to correlate with its activity levels, which may not be necessarily true due to post-transcriptional and translational regulatory processes [9]. To address this issue, a recent direction has been to estimate latent transcription factor activity (TFA) levels from the noisy input prior network, which is also used to constrain the network structure [10–13]. However, the prior knowledge of regulatory interactions is typically noisy, which can significantly influence the quality of the estimated TFA and the subsequent network inference.

Here, we present MERLIN+P+TFA, a regulatory network inference algorithm that extends our previous prior-based inference algorithm, MERLIN+Prior (MERLIN+P) [7], by incorporating a sparsity inducing approach to estimate “regularized TFA” that is robust to noisy input networks. MERLIN+P+TFA extends the Network Component Analysis (NCA, [14]) method to incorporate uncertainty in the input prior networks to enable data-driven selection of the most reliable edges in the TFA estimation. To demonstrate the broad applicability of MERLIN+P+TFA to mammalian species, we collected a large compendium of expression data spanning four commonly used mammalian cell lines, and inferred TFA and regulatory networks. Based on extensive empirical analyses on both simulated and real expression data from yeast, human and mouse species, we show that such estimated TFA is more robust to noise, is more accurate and infers better networks compared to an approach that naively uses the noisy input network. Furthermore, the estimated TFA values are beneficial for other popular expression-based network inference algorithms included several that have previously not been tested with TFA for both bulk and single cell expression datasets. In both human and mouse systems, regularized TFA improves the quality of inferred networks and prioritizes novel regulators that could not be found using expression alone.

We experimentally validated 58 prioritized regulators for the mouse embryonic stem cell (mESC) state using knockdown experiments and phenotypic assays and recover known and novel mESC regulators. We functionally test the predicted target genes of 8 TFs using RT-pCR and find that many of the predicted targets have a significant change in expression indicative of a functional target. Taken together, MERLIN+P+TFA is a powerful tool for reconstructing gene regulatory networks and is broadly useful across diverse systems for both bulk and single cell gene expression datasets.

## Results

### MERLIN+P+TFA: inferring genome-scale GRNs with regularized transcription factor activity (TFA) levels

MERLIN+P+TFA combines expression-based gene regulatory network (GRN) inference with TFA estimation to build a genome-scale regulatory network. The MERLIN+P+TFA algorithm takes as input the gene expression matrix, a noisy prior network and a list of candidate regulator, and performs network inference in two steps (**Figure 1**): 1) estimate TFA using a new approach of NCA, NCA-LASSO, based on regularized network component analysis (NCA), and 2) infer gene regulatory connections using MERLIN+P, a previously developed method, which combines gene expression with prior information [7] (**Section Materials and methods**). MERLIN+P and MERLIN+P+TFA use a “dependency network” [15], a type of “probabilistic graphical model” (PGM) [16], to represent the statistical dependencies between regulators and target genes. The prior network represents putative regulatory relationships between regulators and genes and is typically “noisy” due to both false and missing edges. Given the noisy prior network and expression matrix, the regularized NCA step iteratively estimates the TFA and an updated connectivity matrix. The original NCA algorithm estimated TFA using a pseudo inverse operation, while the connectivity matrix is updated in turn by a per-gene neighborhood selection from a set of candidate regulators from the prior network. In our regularized version, NCA-LASSO, we leverage the confidence of the edge between the regulator and target gene within an L1-based regression framework to constrain the neighborhood selection (**Section Materials and methods**). MERLIN+P takes the estimated TFA matrix and the gene expression matrix and infers a set of regulators for each gene in the dataset. Additionally, MERLIN+P+TFA identifies gene modules where the genes having similar regulatory programs and expression levels are assigned to the same gene module. MERLIN+P+TFA is applied in a stability selection mode to estimate edge and module membership confidence.

**Figure 1.**
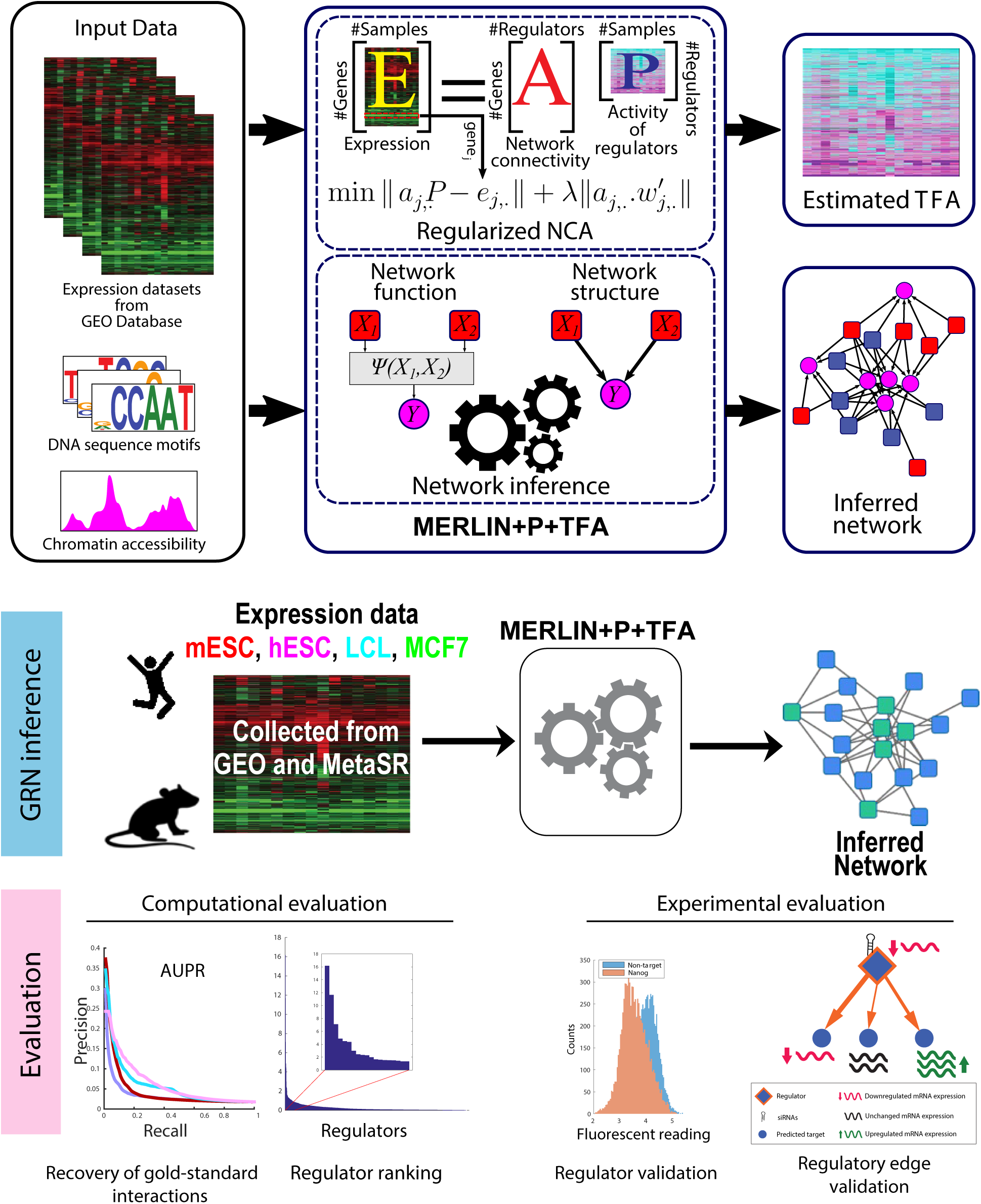
The MERLIN+P+TFA framework for inferring genome-scale gene regulatory networks (GRNs) with hidden transcription factor activity (TFA) levels. MERLIN+P+TFA has two steps: (1) Regularized network component analysis (NCA), (2) GRN inference with MERLIN+P which uses prior knowledge and mRNA and TFA of regulators. Regularized NCA extends the existing NCA method to incorporate the confidence levels in the input network for TFA estimation. This step takes an *m* × *s* genome-scale gene expression matrix, *E* of *m* and a noisy prior network connecting the genes to *p* regulators, as input and estimates the TFA *P* of the *p* regulators by minimizing the error ||*E* − *AP* ||^2^. Here, “*A*” is the updated “connectivity matrix” of the regulators and target genes and is initialized based on the input prior network. *A* and *P* are estimated using an iterative algorithmy. In regularized NCA, the objective is extended to add a regularization accounting for the confidence of a regulator-target pair (*w_ji_*) available in the prior. The output of MERLIN+P+TFA is the TFA levels and the final inferred network. MERLIN+P+TFA was applied to simulated and real expression datasets from yeast, human and mouse. The inferred GRNs were evaluated using multiple metrics (AUPR, predictable TFs, regulator ranking). The mouse embryonic stem cell (mESC) network was further used to nominate ESC-specific regulators and experimentlaly validated with regulator knockdown expreriments and target expression post regulator knockdown.

### Regularized NCA accurately estimates TFA and GRNs with simulated data

We first assessed the performance of MERLIN+P+TFA’s regularized NCA estimation algorithm, NCA+LASSO and downstream network inference when considering input prior networks with different extents of noise.

To this end, we performed a simulation experiment with ground truth TFA and network structures, which were used to simulate an expression dataset (**Section Simulation experiments**). The noisy prior networks were created by randomly adding and removing edges from the true network and assigning edge weights using two different beta distribution whose parameters, *a* and *b* can be used to change the skew of the distribution (**Figure S1**). In the “uniform” setting (*a* = 1, *b* = 1), the edge weights of the true and false edges were set at random, and did not discriminate the true/false status of the edges. In the “non-uniform” setting (*a* = 5, *b* = 1), the true edges tend to have higher edge confidence than the false edges. For each of the edge weight schemes, we prepared three versions of the prior network by introducing 10%, 30%, and 50% noise by randomly adding or removing edges. The simulated expression dataset along with the prior network were used to estimate the TFA. Subsequently, we used the estimated TFA, the prior network, and the simulated expression dataset to infer a network using the MERLIN-P algorithm. This allows us to compare the estimated TFA to true TFA, and inferred network to true network.

We first compared the estimated TFA using the original NCA algorithm from Liao et al [14], and true TFA across different noisy edge and edge weight settings. As expected, when we increase the noise level in the input prior network, the agreement between the estimated TFA and true TFA decreases irrespective of the edge weight scheme (**Figure 2A, B**). For example, the median correlation decreases from 0.79 to 0.57 to 0.44 when the noise level increases from 10% to 30% to 50% in the case of uniform edge weights (**Figure 2A**). We also observed that the networks inferred from the estimated TFA have lower agreement (as measured by AUPR) to the true network as we increase the noise, again irrespective of the edge weight scheme (**Figure 2B**). When using non-uniform edge weights in the prior, the AUPR values at the same noise level are higher than that with uniform edge weights. This is because MERLIN+P incorporates the edge weight information in the inference process [7]; hence, a prior network with more accurate edge weights results in a more accurate inferred network, while a more noisy prior network produces an inferred network of a lower quality and lower AUPR.

**Figure 2.**
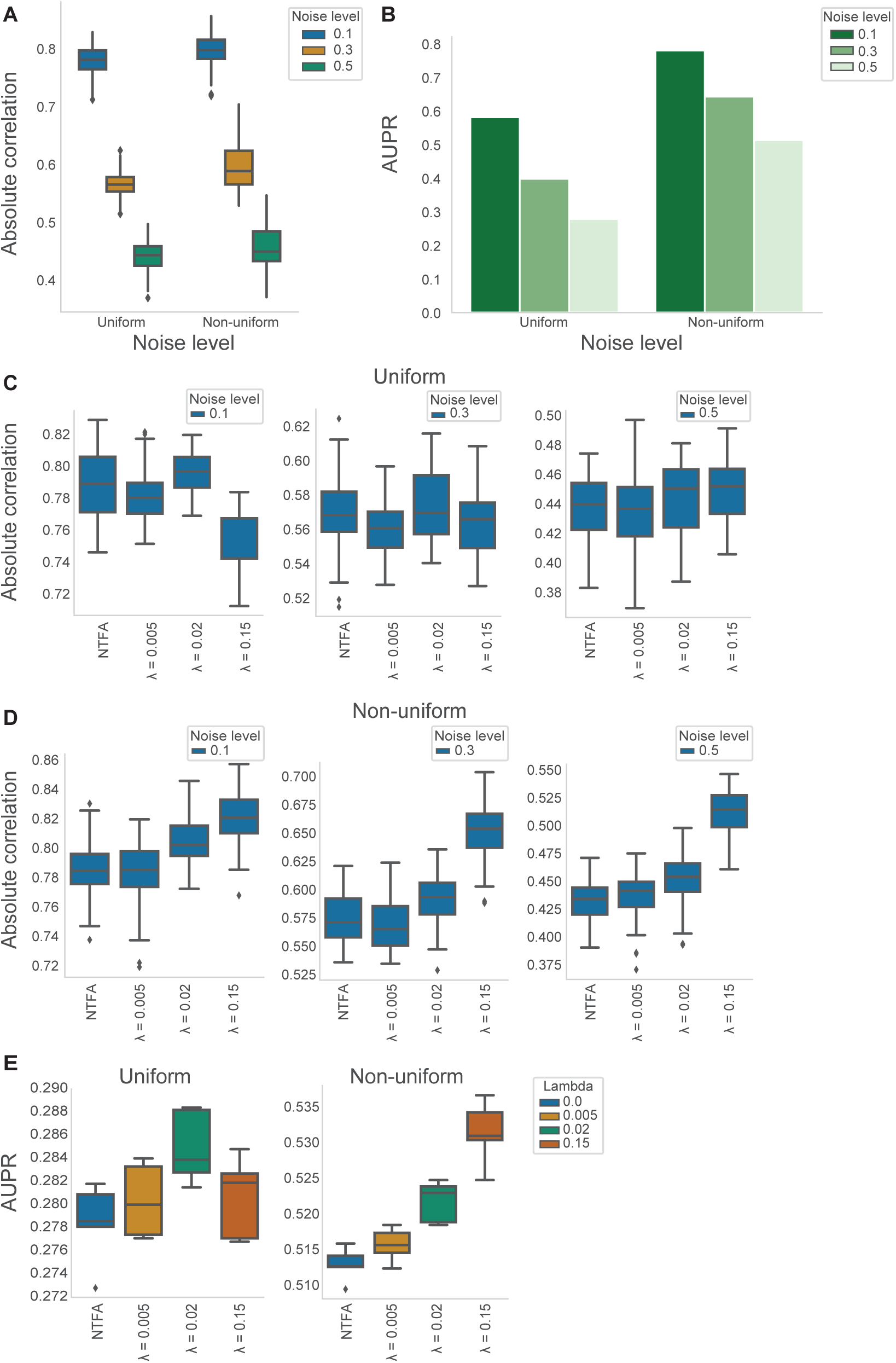
The effects of the prior network properties on the performance of NCA and regularized NCA with the simulated data. “Non-uniform” refers to the (*a* = 5, *b* = 1) configuration. (**A-B**) The effect of including noise in the prior network on NCA’s performance. (**A**) The absolute correlation between the TFA estimated from a noisy prior network (with 10%, 30%, or 50% noise) and the true TFA. (**B**) The area under the precision-recall curve (AUPR) values of the network inferred from the TFA estimated using the noisy prior network at a specific noise level. (**C-E**) The comparative performance of NCA and the regularized NCA (NCA+LASSO) in (**C-D**) estimating TFA for absolute correlation with 10%, 30%, or 50% noise and (**E**) inferring network given the prior network with 50% noise.

Next, we asked whether incorporating prior edge weights through regularization can improve the quality of the estimated TFA. We used the prior network with 10%, 30%, and 50% noise and estimated TFA using NCA+LASSO at different regularization levels (**Section Simulation experiments**), and compared the estimated TFA to the true TFA based on absolute correlation (**Figure 2C, D**). For both uniform and non-uniform edge weight distributions, the correlation between the true and NCA-LASSO estimated TFA improve and are higher than the correlation when using TFA estimated using unregularized NCA (NTFA, **Figure 2C, D**). When the edge weights follow the non-uniform distribution (**Figure 2D**), regularization significantly improves the correlation between the true TFA and estimated TFA. Finally, the AUPR values of the networks inferred from the NCA+LASSO-based TFA are higher than networks inferred using NTFA (**Figure 2E**), the differences in AUPRs being more striking for the non-uniform edge weights. Taken together these results suggest that regularized NCA approach, NCA-LASSO, effectively incorporates noisy prior networks providing more accurate estimations of TF activity and GRNs than the unregularized version, which is more susceptible to increasing noise in the prior network.

### Regularized TFA is beneficial for diverse network inference methods

We next investigated the performance of regularized TFA of NCA-LASSO on a real dataset from the yeast, *Saccharomyces cerevisiae*. We used a motif-based network as input prior and expression from a large scale natural variation dataset (**Section Materials and methods**). We estimated TFAs and downstream regulatory networks using both regularized and unregularized versions of the NCA algorithm. For network inference, in addition to MERLIN+P, we included several other network inference methods, namely GENIE3 [17], Inferelator [6, 18], TIGRESS [19], mLASSO-StARS [13], NetREX [20], MERLIN [21], and MERLIN+P [7]. Among these methods, MERLIN+P+TFA, mLASSO-StARS, Inferelator, and NetREX explicitly infer TFA and utilize the estimated TFAs for network inference. MERLIN+P+TFA, MERLIN+P, Inferelator, etREX, and mLASSO-StARS also incorporate the prior network during network inference in addition to using the prior network for TFA estimation. We applied these algorithms with expression alone (no TFA), with unregularized TFA based on Network Components analysis (NTFA) and regularized TFA (RegTFA) (**Table S1**).

We evaluated the inferred networks based on AUPR scores with respect to three gold standard networks (**Section Expression datasets**), regulator ChIP binding (ChIP), regulator knock out or knock down (KD), and the intersection of the ChIP and KD gold standard networks When an algorithm had hyper-parameters (e.g., regularization of TFA), we used the setting with the highest AUPR for that algorithm (**File S1**, **File S2**). MERLIN+P+TFA outperformed all other algorithms across the three gold standard networks when comparing methods that use expression alone, expression with prior, or unregularized TFA (**Figure 3A**). Interestingly, methods that explicitly incorporate unregularized TFA do not always outperform methods that do not (**Figure 3A** gold bars). When comparing networks inferred without TFA, (**Figure 3A** blue bars), incorporation of prior (MERLIN+P) is better than not using prior for all three gold standard networks. We next asked whether regularized TFA is beneficial for methods other than MERLIN-P including methods like GENIE3 and TIGRESS which were developed for expression alone. We find that compared to not using TFA or unregularized TFA, incorporation of TFA is beneficial for network inference for all methods compared (**Figure 3B**, green vs gold bars).

**Figure 3.**
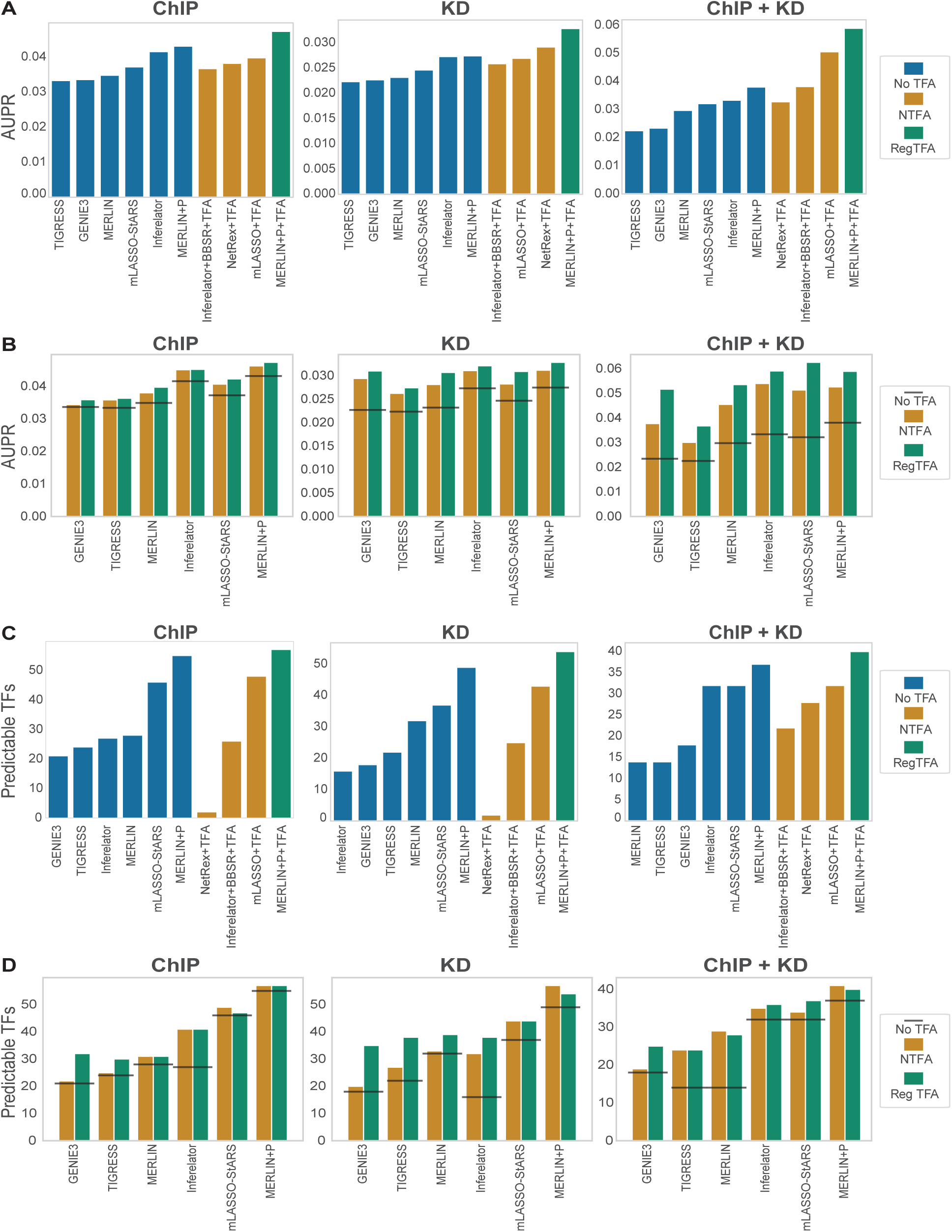
Performance of GRN inference methods on the yeast dataset. (**A**) AUPR scores of methods that were applied without including TFA (No TFA), with unregularized TFA (NTFA), or using regularized TFA (RegTFA) with respect to the ChIP, KD, and the intersection of the ChIP and KD gold standard networks (ChIP + KD). Shown are the highest AUPRs across different regularizations. For {GENIE3, TIGRESS, MERLIN, MERLIN+P}, the unregularized AUPR scores correspond to the TFA estimated with NCA (*λ* = 0.000). For {mLASSO-StARS, Inferelator}, the unregularized AUPR scores correspond to the TFA internally estimated by the methods themselves. We exclude method NetREX since it could not be used without its internally estimated TFA. (**B**) AUPR scores of the inferred networks comparing regularized (RegTFA) and unregularized TFA (NTFA) across the three gold standards. No TFA is shown as a horizontal line for the respective algorithm. (**C**) The number of predictable TFs in the inferred networks. Shown are the highest value of this metric across different regularizations computed using top 30k edges. (**D**) Predictable TFs of inferred networks for regularized (RegTFA) and unregularized TFA (NTFA) across the three gold standards. The horizontal line depicts the No TFA setting.

As AUPR is a coarse metric for assessing the entire inferred network structure, we next examined recovery of targets of individual TFs using the “predictable TFs” metric. This is defined as the number of TFs whose targets in the inferred network have a significant overlap with their targets in the gold standard network (**Section Additional evaluation metrics for real expression datasets**). To do so, we counted the number of “predictable TFs” in each inferred network with respect to a given gold standard network with different TFA settings. Across all methods compared, the highest number of predictable TFs is achieved when they utilized prior or TFA or both along with the expression data (**Figure 3C**). When comparing regularized vs unregularized TFA, most methods either benefit or do not suffer in performance (**Figure 3D**). Exceptions to this were mLASSO-StARS on ChIP and MERLIN+P+TFA on KD and ChIP+KD, which exhibited a small reduction in performance. We also examined the enrichment metric for each TF and found that the overlap between its true and predicted targets tends to be more significant when TFA is utilized along with the expression data (**Figure 4**). Overall, we observed that the addition of TFA improves the number of predictable TFs, and regularized TFA performs better than original TFA in almost all the cases (11 out of 16 cases while having ties in 2 cases).

**Figure 4.**
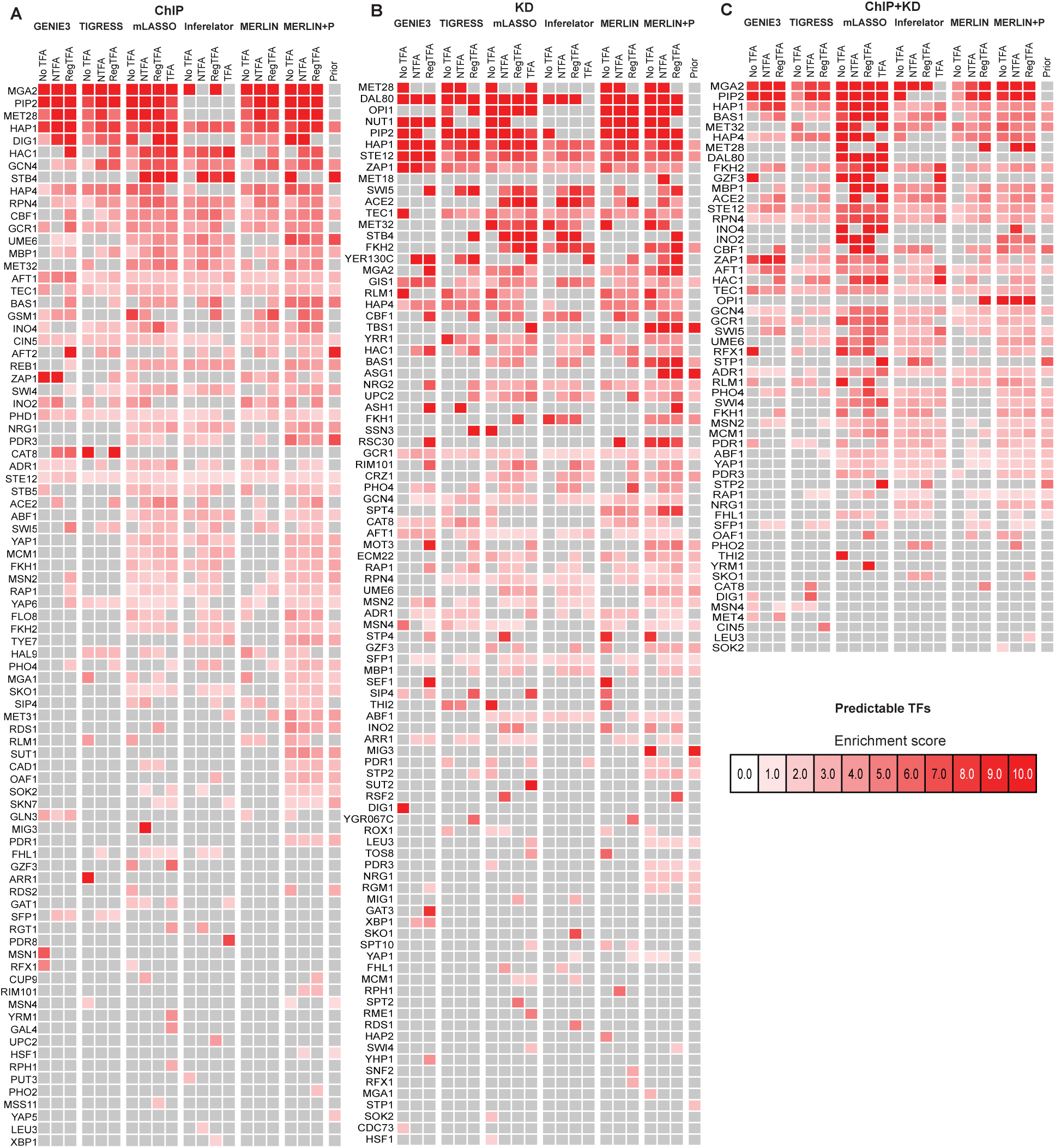
The enrichment scores of each TF (row) in the networks inferred by different algorithms for different TFA settings (NTFA, RegTFA, TFA) and different gold standard networks–(**A**) ChIP, (**B**) KD and (**C**) Intersection of the ChIP and KD (ChIP+KD). NTFA: no regularization; RegTFA: regularized TFA with the best metric; TFA: shown for mLASSO-StARS and Inferelator represents the TFA estimated by each algorithm’s inbuilt TFA estimation process. The higher the enrichment score, the more significant the overlap is between the targets of a TF between the inferred and gold standard network.

### MERLIN+P+TFA for inferring genome-scale transcriptional regulatory networks in mammalian cell lines

Having demonstrated the benefits of MERLIN+P+TFA’s regularized TFA in yeast, we next applied it to mammalian datasets from four different cell lines: the mouse embryonic stem cell (mESC) line, human ESC (hESC) line, human Lymphoblastoid cell line (LCL), and human Michigan Cancer Foundation-7 cell line (MCF7). For each of the cell lines we collected a large dataset of published RNA-seq expression samples in mouse embryonic stem cell (**Section Expression datasets**). For mESC, we additionally collected a microarray dataset resulting in a total of 5 datasets. For each of the datasets, we used a motif-based prior network filtered by DNase I footprints, and estimated TFA at different levels of regularization (**Section MERLIN+P+TFA for GRN inference from bulk gene expression data**). We applied MERLIN and MERLIN-P with expression alone (no TFA), with unregularized TFA based on Network Components analysis (NTFA) and regularized TFA (RegTFA). We compared the inferred networks based on AUPR, and predictable TFs across three gold standards for each cell line and dataset (**Figure 5**). For networks inferred using RegTFA, we selected the regularization setting that returned the highest AUPR for further comparisons (**File S3**, **S4**). Based on global AUPRs (**Figure 5**), both MERLIN and MERLIN-P had higher AUPRs when using TFA versus not and especially with RegTFA. Furthermore, MERLIN+P obtained higher AUPR scores than MERLIN; further demonstrating the utility of using priors for network inference. Across the three types of gold standards, addition of prior was much more useful on ChIP and ChIP-KO gold standards.

**Figure 5.**
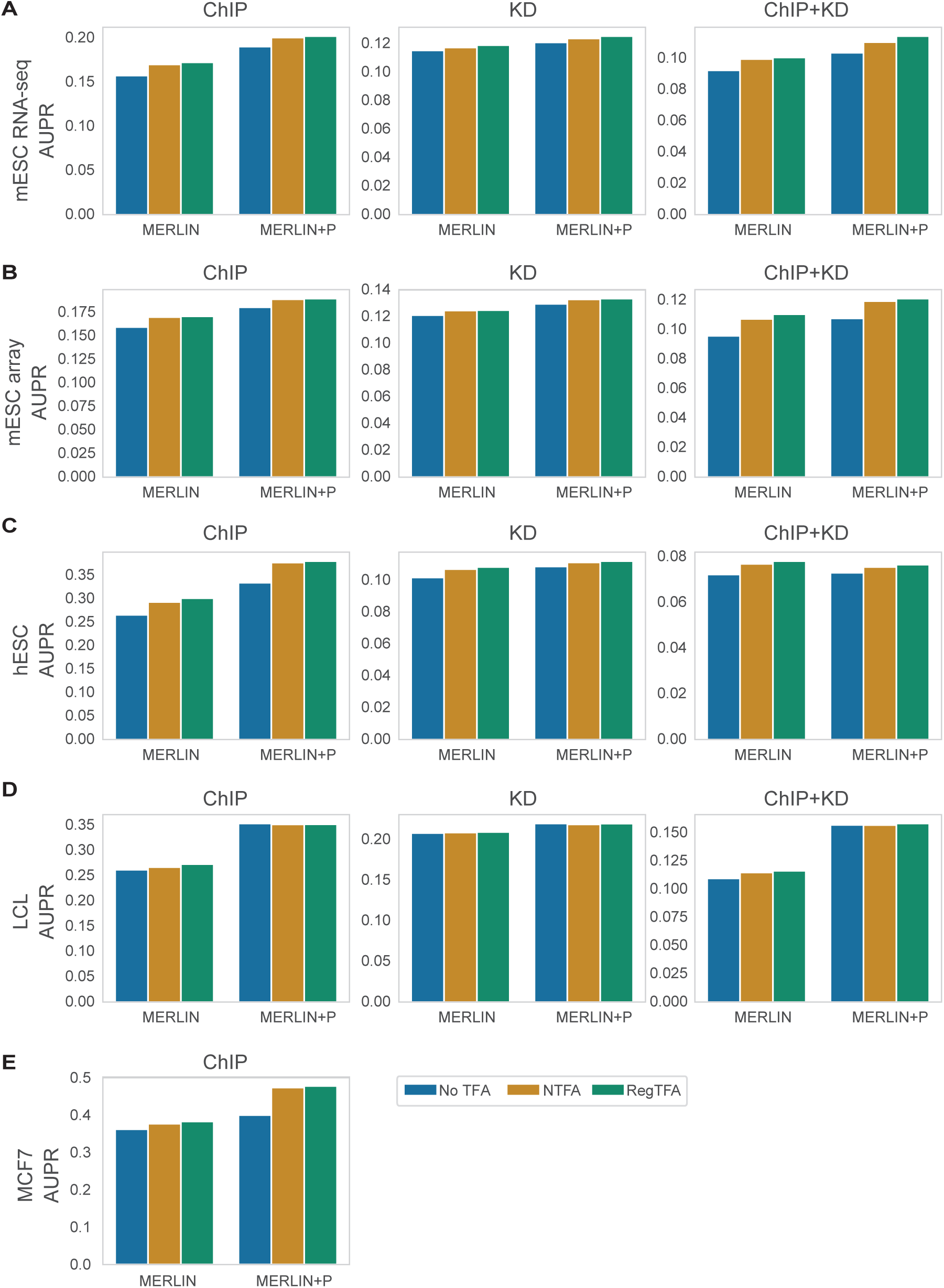
The AUPR scores of the networks inferred with MERLIN and MERLIN+P from five mammalian datasets: mESC RNA-seq (**A**), mESC array (**B**), hESC (**C**), LCL (**D**), MCF7 (**E**), on three gold-standard networks (ChIP, KD, and the intersection of ChIP and KD (ChIP+KD)). Shown are AUPR scores for each method for three TFA settings: expression data only (No TFA), NCA having *λ* = 0 (NTFA), and regularized TFA (RegTFA). The best performing regularized NCA result was selected for each comparison and is represented as “RegTFA”.

We next examined the per TF recovery of the structure based on the predictable TF metric. MERLIN achieved the highest number of predictable TFs for regularized TFA for 4 out of the 13 cases (each case corresponding to a dataset-gold standard combination) and was higher compared to expression alone for 6 out of 13 cases (**Figure 6**). On the other hand, MERLIN+P achieved its highest numbers of predictable TFs with regularized TFA for 5 out of 13 comparisons. With respect to comparing MERLIN to MERLIN+P per TFA treatment for the ChIP gold standard, MERLIN+P always predicted more predictable TFs than that of MERLIN across all datasets, except for expression alone and regularized TFA for hESC (**Figure 6C**). We observe a similar behavior in the other cell lines as well for the other gold standards and datasets as well (**Figure S2-S6**). The exceptions to this are a few KD based comparisons of the mESC array dataset (**Figure 6B**) and hESC dataset where MERLIN with TFA was worse (**Figure 6C**). For the ChIP gold standard intersected with the KD gold standard, MERLIN+P performed better than MERLIN for mESC RNA-seq and the LCL cell line, but this pattern was not consistently replicated for the mESC array and hESC dataset applications (**Figure 6A-C**). Overall, for the mammalian cell lines, the predictable TFs metric was best when using regularized TFA demonstrating the advantage of using MERLIN+P+TFA for mammalian datasets.

**Figure 6.**
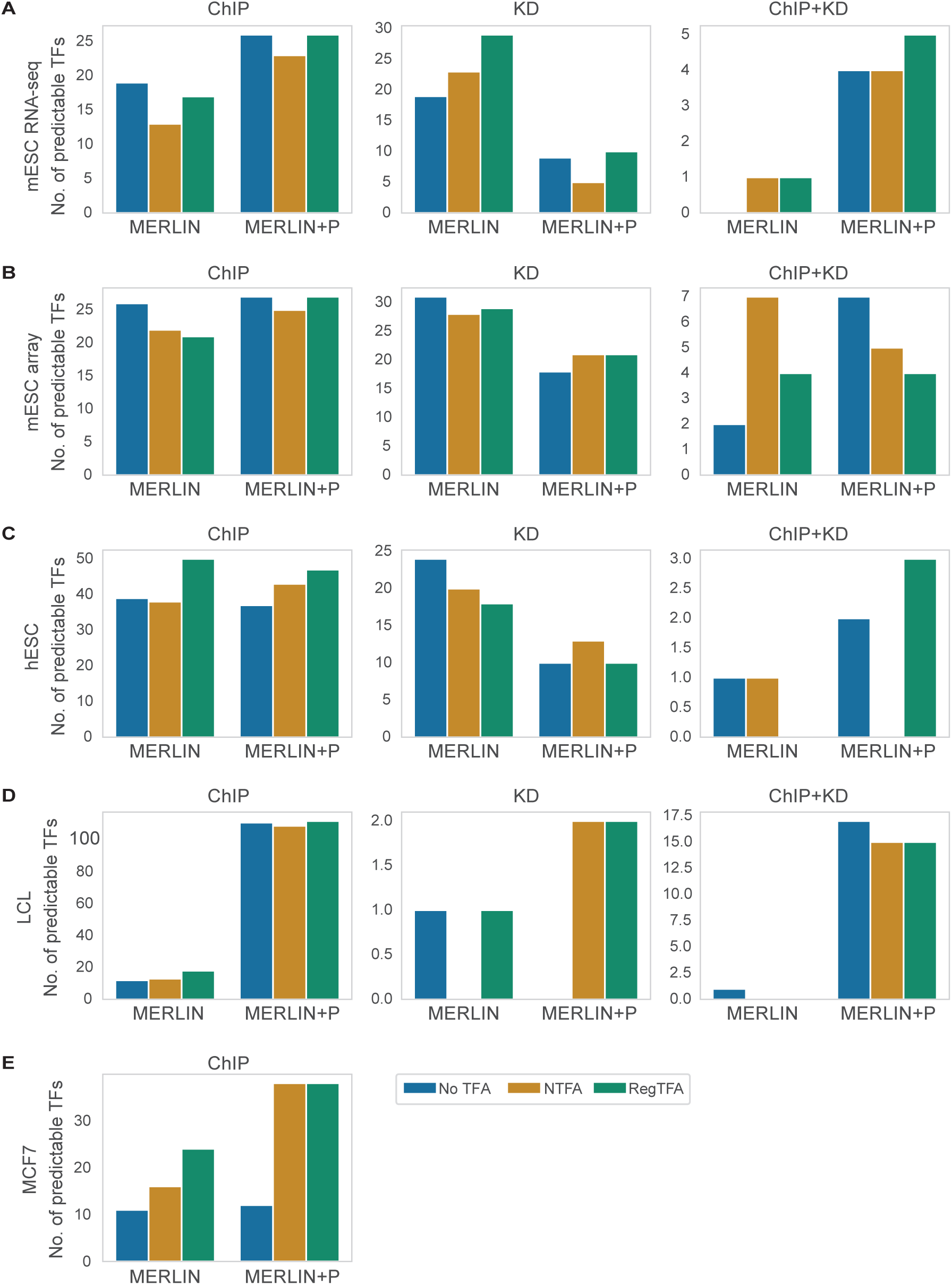
Number of predictable TFs in the networks inferred with MERLIN and MERLIN-P from five mammalian datasets, mESC RNA-seq (**A**), mESC array (**B**), hESC (**C**), LCL (**D**), MCF7 (**E**) on three gold-standard networks (ChIP, KD, and the intersection of ChIP and KD (ChIP+KD)). Shown are predictable TF counts for each method for three TFA settings: expression data only (No TFA), NCA having *λ* = 0 (NTFA), and regularized TFA (RegTFA). The best performing regularized NCA result was selected for each comparison and is represented as “RegTFA”. Predictable TFs are computed on the top 30k edges.

### Experimental validation experiments for the mESC transcriptional regulatory network

Given MERLIN+P+TFA’s good performance in both mammalian and yeast datasets, we used its predictions to find new regulators and targets important for the mouse ESC state. We first ranked regulators by an importance metric that assessed the change in predictive power of the inferred network if the regulator was removed from the network (**Section Additional evaluation metrics for real expression datasets**). We considered both array and RNA-seq-based networks inferred using both MERLIN and MERLIN-P (**Figure 7A**). We evaluated this ranking based on the position of genes known to be important for the ESC state. To create the list of known ESC genes, we combined experimentally derived gene sets from Wong et al. [22] and Mueller et al. [23] available from MsigDB [24, 25] together with genes annotated with ESC-relevant Gene Ontology (GO) terms [26], “regulation of embryonic development”, “embryonic organ development”. MERLIN+P+TFA achieved the highest AUPR scores with regularized TFA for both array and RNA-seq networks. This pattern is true for MERLIN as well indicating the importance of regularized TFA for prioritizing important cell-state specific regulators (see **File S5** for gene module figures) . Among the highly ranked genes were Esrrb and Nanog (**Figure 7B**, **Figure S7**) that are key components of the pluripotency GRN.

**Figure 7.**
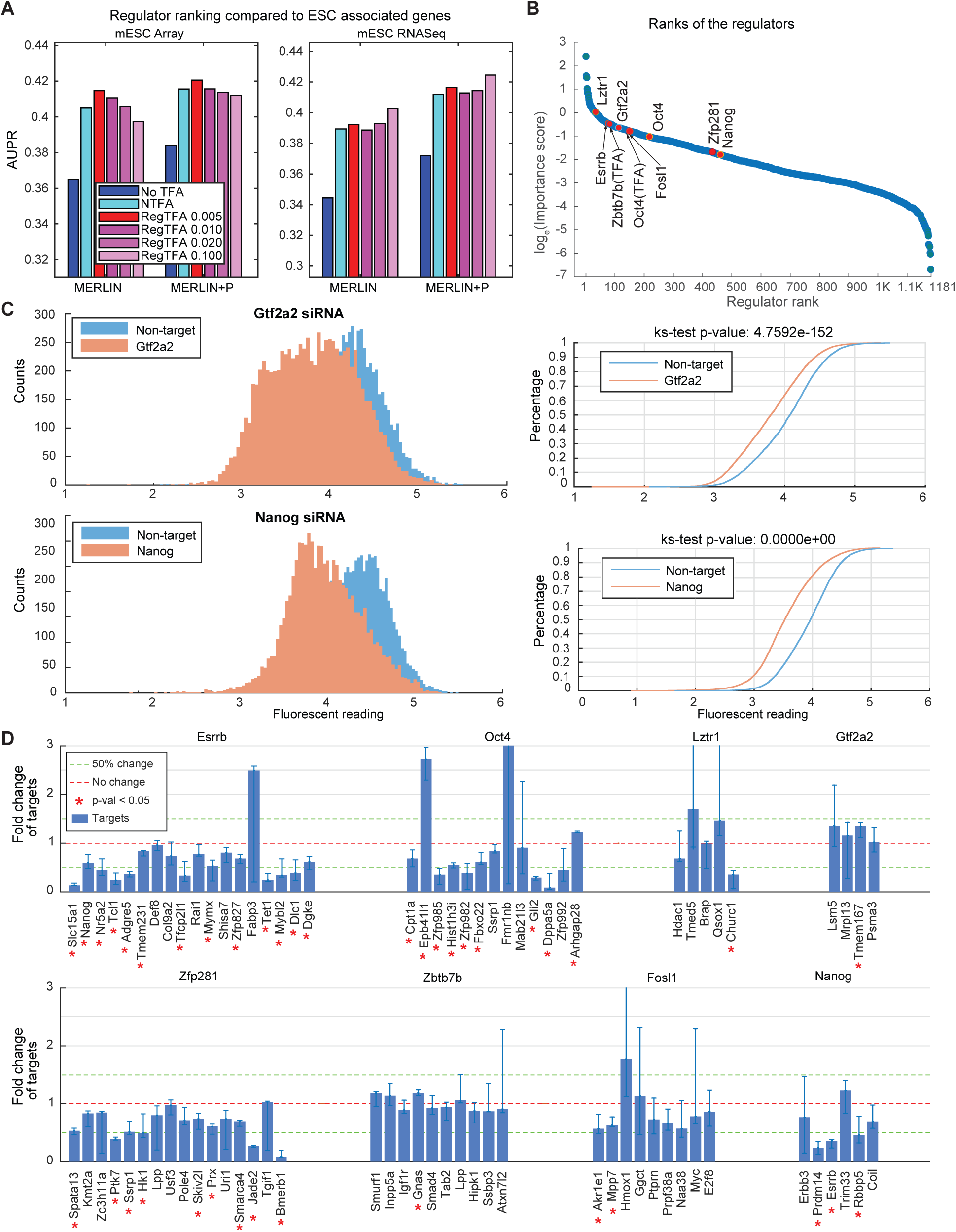
Experimental validation of the inferred mESC GRN. (**A**) The AUPR of the top *N* regulators in each inferred network of {MERLIN, MERLIN-P} for six different TFA settings (No TFA, NTFA, RegTFA with different regularizations) on the mESC array and RNA-seq datasets. The AUPR was computed using a gold-standard list of ESC related genes curated from the literature. (**B**) Regulators (1s, 181) in the MERLIN+P+TFA inferred network ranked based on importance score. Shown are rankings using regularized TFA *λ* = 0.01 (TFA0.010) from the mESC RNA-seq dataset. Four known ESC regulators (Esrrb, Nanog, Oct4, Zfp281) and 4 novel (Fosl1, Gtf2a2, Zbtb7b, Lztr1) TFs are highlighted in red cicles. “Oct4” is highlighted twice since it in the inferred GRN using gene expression (Oct4) and its estimated TFA (Oct4 (TFA)). (**C**) The effect of the siRNA knockdowns of “Gtf2a2” (a novel TF) and “Nanog” (a known TF). Left: histograms showing the distributions of Nanog fluorescence for Gtf2a2 (top) and Nanog (bottom) siRNA knockdown cells in the E14 cell line. The higher the intensity, the higher the protein expression of Nanog per cell, which is used as a proxy for measuring cellular pluripotency. The orange distribution is for the TF knockdown and the blue distribution is for “non-targeting siRNA control” experiments. For both Gtf2a2 and Nanog, the knockdown distributions (orange) is shifted significantly to the left with respect to the control distributions (KS test p-value *<*1E-10). Right: cumulative distribution function (CDF) of the respective histograms on the left. (**D**) Fold change of expression of targets on knocking down one of the 8 TFs as measured by RT-PCR. The targets with statistically significant fold change (t-test *p*-value*<* 0.05) are marked with asterisks. Please note that only the top 50, 000 edges of each inferred network were considered for generating the panels in this figure.

We next used these rankings to identify novel regulators important for maintenance of the ESC state and also to validate targets of selected regulators. Briefly, we took the union of the top 10% of regulators in all 10 MERLIN+P+TFA networks (both array and RNA-seq, for all TFA regularization settings i.e. all settings excluding “No TFA”). The union resulted in a list of 205 regulators. We examined the literature for these regulators and excluded 147 regulators already most of which are known to be important for the maintenance of or differentiation from the ESC state (**Table S2**). We kept a few known regulates, e.g., Esrrb, Zfp281, as positive controls. These remaining 58 regulators, the majority of which have previously not been associated with pluripotency, were tested with small interfering RNA (siRNA) knockdown experiments (**Section Experimental validation of MERLIN+P+TFA mESC GRN**). Briefly, we used siRNAs to knock down each of the regulators at a time and measured the expression of Nanog, a marker of pluripotency. We compared the distribution of Nanog in the regulator-targeted cell population and a control population targeted by a control siRNA using the Kolmogorov-Smirnov (KS) test (**Table 1**, **File S6**). We did these experiments in the 2D4 and E14 ESC lines. Nanog’s expression was measured with the Nanog-GFP reporter in the 2D4 cell line and through intracellular staining in the E14 cell line. Out of these 58 novel regulators, 19 regulators showed significant decrease in pluripotency of the 2D4 cell line and 18 showed significant decrease in the E14 cell line (FDR corrected *p*-value*<*1E-10). Among these regulators, 10 are common in both the cell lines. There are 11 and 23 TFs in the 2D4 and E14 cell lines, respectively, that show significant increase (*p*-value*<*1E-10) in pluripotency as a result of knock down (7 of them are common in both the cell lines).

We next tested the targets corresponding to the high-confidence edges in the MERLIN+P+TFA inferred network of 4 novel TFs that are not present in the mESC gold standard networks (namely, Fosl1, Gtf2a2, Zbtb7b, Lztr1) and 4 known TFs (Esrrb, Nanog, Oct4, Zfp281), by measuring the expression of the targets (with RT-PCR) after the siRNA knockdowns of these TFs. We examined the fold changes in the expression levels of the targets compared to that of the non-target negative controls (Kdm3b, Cbx3, and AF9, dashed red line **Figure 7D**). 39 out of the 81 tested TF-target pairs (48%) show significant change in expression compared to control. Of these 39 pairs, 5 correspond to the targets of the 4 novel TFs. The remaining 34 pairs correspond to the targets of the known TFs. Among them, 24 TF-target pairs were already part of the ChIP and KD gold standard networks (6 pairs belonging to ChIP only, 5 pairs belonging to KD only, and 13 pairs belonging to both ChIP and KD; see **File S7**). Subsequently, we investigated how many of these 39 TF-target pairs had been predicted using the estimated TFA profiles of the regulators. It was observed that 20 out of 39 pairs had been predicted using the estimated TFA profiles of the corresponding regulators while the remaining 19 pairs had been predicted using the gene expression levels of the regulators; no pairs had been predicted using both the TFA profiles and gene expression levels of the regulators (see **File S7**, **S8**). Thus, both expression and TFA are important in the identification of important regulators and MERLIN+P+TFA was able to effectively combine both modalities to infer more complete and accurate estimation of GRN structure. Taken together, these experiments suggest that our predicted regulators and high-confidence inferred interactions can provide useful insight into the regulatory mechanism of pluripotency.

### Regularized TFA improves network structure recovery for scRNAseq datasets

The availability of single-cell RNA sequencing (scRNAseq) datasets opens up new opportunities for genomescale GRN inference by leveraging the variation in expression across individual cells. The effect of TFA incorporation for learning GRNs on scRNAseq datasets has been examined on a limited scale [27–29], and has shown that TFA incorporation often increases network performance. Given the advantage of MERLIN+P+TFA’s regularized TFA on bulk gene expression data, we next asked if regularized TFA is beneficial for scRNA-seq GRN inference as well. We selected three network inference algorithms, MERLIN-P [7], Inferelator [30], and Single-Cell rEgulatory Network Inference and Clustering (SCENIC) [31] to compare across six scRNA-seq. We selected these algorithms because these were among the top performing methods in our previous benchmarking study of scRNA-seq network inference methods [29], and allowed us to extend the scope of previous findings here with the addition of regularized TFA. Additionally, MERLIN-P and Inferelator incorporated priors for GRN inference, which we could utilize for comparisons. We obtained datasets from McCalla *et al.* [29], which encompassed samples from three species, yeast, mouse, and human, and represented dataset sizes of varying numbers of cells (**Materials and methods**), ranging from 163 cells [32] to 17,396 cells [30]. As before, we used the Perturb, ChIP, and the intersection of Perturb and ChIP gold standards and AUPR, F-score and predictable TFs as metrics, to evaluate the performance of each algorithm. When networks were inferred for an algorithm with multiple regularized TFA settings, the setting that returned the highest AUPR was used for comparisons (**File S9, S10**).

Based on F-score across all three types of gold standards, regularized TFA (RegTFA for non-MERLIN algorithms) or just TFA for MERLIN+P+TFA) was generally better or at par than unregularized TFA estimated by the algorithm (**Figure 8A**, **S8**, **File S10**). For Inferelator the performance depended on how the unregularized TFA was estimated, with regularized TFA (Inferelator+P+RegTFA) being better than Inferelator’s unregularized TFA (Inferelator+P+TFA), but being outperformed when using the NCA algorithm (Inferelator+P+NTFA). We observed a similar behavior when using AUPR and predictable TFs, with a greater benefit when using ChIP-based gold standards.

**Figure 8.**
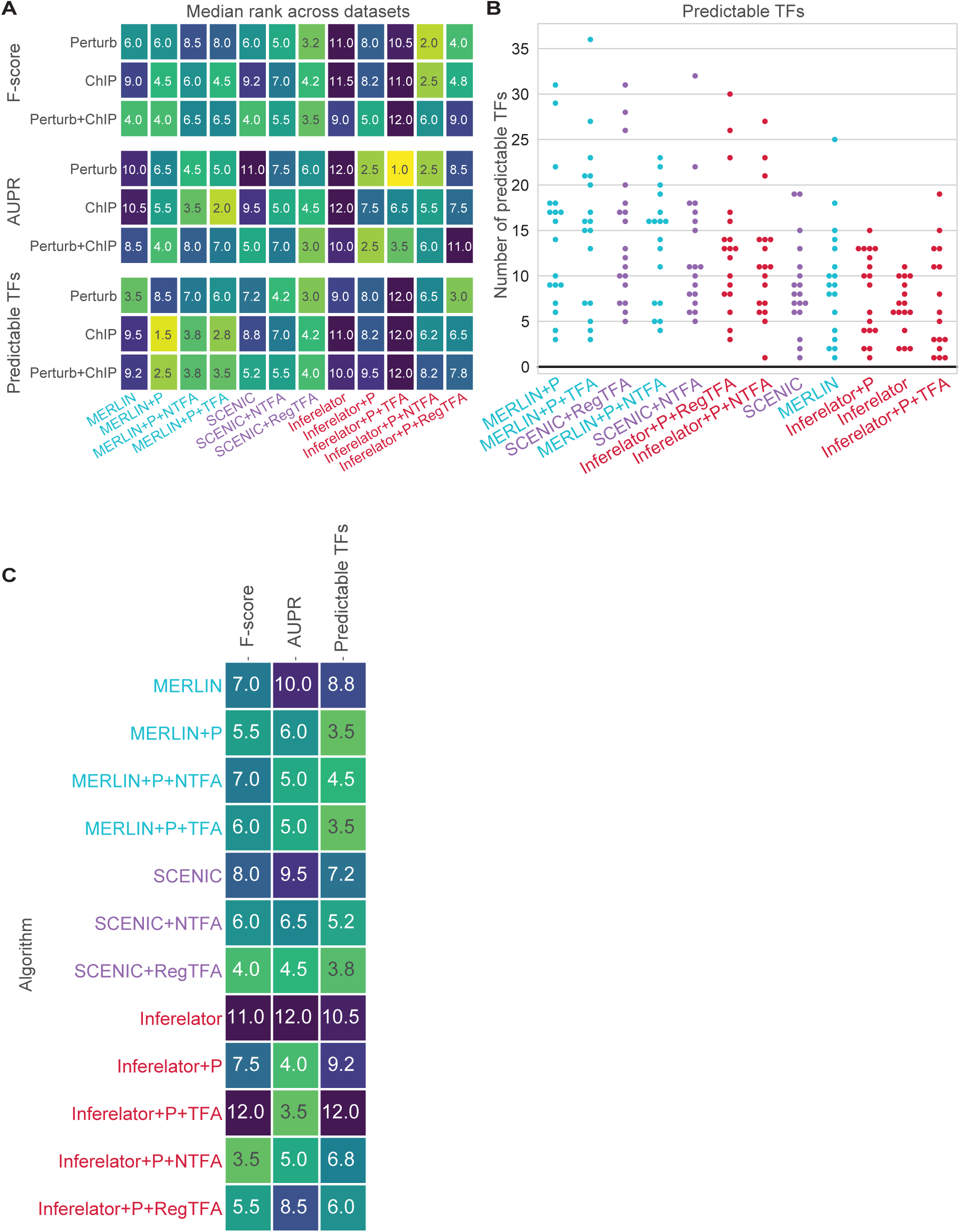
Assessing regularized TFA for GRN inference on scRNAseq datasets.(**A**) Algorithm median rank across 6 scRNAseq datasets based on performance on ChIP, Perturb, and intersection of the ChIP and Perturb (Perturb+ChIP) gold standards. Performance is measured with F-score, AUPR, and predictable TF metrics. (**B**) Predictable TF distribution for all scRNAseq datasets (Gasch, Jackson, A2S, FBS, Shalek, Han) and gold standard networks (ChIP, Perturb, Perturb+ChIP). Algorithms are ordered according to median rank performance (top performing algorithm placed on the left and the bottom performing algorithm placed to the right). (**C**) Overall median algorithm ranking across gold standards and scRNAseq datasets. Algorithm ranking is shown for F-score, AUPR, and predictable TFs metrics.

When comparing within one algorithm across different TFA settings, GRN inference generally improved in quality when using TFA versus not, but it depended on the algorithm. For example, SCENIC+RegTFA outperformed SCENIC and SCENIC+NTFA across all three gold standard networks for F-score (**Figure 8A**, **S8**, **File S10**). MERLIN+P+NTFA or MERLIN+P+TFA outperformed MERLIN alone 5 out of 9 times across metrics and gold standards. The difference between using Prior (MERLIN+P) versus TFA was less clear for MERLIN, especially when using Perturb or Perturb+ChIP. Similarly, Inferelator with Prior (Inferelator+P) was ranked highest on the Perturb+ChIP gold standard. We observed a similar trend when using predictable TFs and AUPR. However when performance was combined across all of the gold standards, Inferelator+P+RegTFA outperformed all of the other TFA configurations for Inferelator (**Figure 8B, C**).

Overall trends summarized across gold standards and metrics show algorithm configurations that include RegTFA are among the top-ranked algorithms (**Figure 8B, C**). In particular, two of the top 3 ranking algorithms for F-score include TFA for network inference (Inferelator+P+NTFA, SCENIC+RegTFA, Inferelator+P+RegTFA). For AUPR, not only did the top ranked algorithm included TFA (Inferelator+P+TFA), the bottom three ranking algorithms did not incorporate a prior or TFA (MERLIN, SCENIC, Inferelator, **Figure 8B, C**). MERLIN+P+TFA tied for the best ranked algorithm for the predictable TFs metric along with MERLIN+P. In fact, when evaluated for predictable TFs, six out of the seven best performing algorithms included TFA for network inference. Taken together, these results demonstrate that MERLIN+P+TFA’s regularized TFA can benefit network inference for scRNA-seq datasets as well and importantly this benefit is apparent for algorithms like SCENIC which do not have inbuilt capability of estimating TFA.

## Discussion

Reconstruction of genome-scale gene regulatory networks is a long-standing problem in gene regulation and remains experimentally and computationally challenging. Recent computational methods have advanced the field beyond purely expression-based network inference by incorporating network structure priors [6–8, 21, 33, 34]. The prior can not only constrain the inferred networks with biological knowledge, but also estimate the hidden transcription factor (TF) activity levels, which helps relax the assumption that mRNA and protein levels of the TF must correlate [10–13]. However, the prior can be noisy, which can adversely affect the estimated TFA and downstream network inference. To address this problem, we developed MERLIN+P+TFA, that uses regularized regression to selectively incorporate high confidence prior edges enabling more robust TFA estimation and network inference. Beyond MERLIN+P+TFA, the regularized TFA can benefit diverse state-of-the-art network inference methods that may or may not have originally used TFA for both bulk and single cell RNA-seq datasets.

Studies of TFA and GRN inference have thus far been conducted in yeast and bacteria with only a handful in multi-cellular organisms [13, 20]. Our work examined four commonly used mammalian cell lines: mouse embryonic stem cell (mESC), human ESC (hESC), human lymphoblastoid cell line (LCL), and human Michigan Cancer Foundation-7 cell line (MCF7) by systematically collecting expression data, prior information and gold standards in these cell lines. We applied both MERLIN-P and MERLIN+P+TFA to these datasets and showed that regularized TFA improved the quality of inferred networks for these cell lines. Our results substantially increases our understanding of the utility of TFA estimation in mammalian GRNs and are relevant to a large community of researchers. The resources we have created are available to the community to build new inference methods as well as to benchmark them.

Given the increasing availability of single cell RNAseq (scRNAseq) and their potential to capture cell type specific regulation, we studied the effect of regularized TFA for GRN inference from scRNAseq datasets. Across different datasets, we found that regularized TFA generally improved the quality of inferred networks across datasets from different species and algorithms. Our results are consistent and build on previous benchmarking studies of GRN inference from scRNAseq datasets that showed that prior information incorporated as constraints on the GRN structure and/or TFA estimation can improve GRN inference compared to purely expression based GRN inference. Importantly this finding holds for multiple inference algorithms across datasets from different species.

An unmet challenge in the field of GRN inference is the lack of sufficient ground truth regulatory interactions, especially for mammalian systems. The large number of putative regulatory interactions that could be tested experimentally make the generation of comprehensive gold standard datasets expensive, if not, impractical without any guided supervision. Model-guided iterative refinement of GRN structure can make the generation of relevant experimental datasets more efficient. To this end, we used MERLIN+P+TFA network to both prioritize important regulators, as well as select targets of select regulators to validate the regulatory relationships for the mouse embryonic stem cell state (mESC). Among the top 10% regulators ranked by MERLIN+P+TFA, the majority (147 out of 205 regulators) are already known to be associated with pluripotency, indicating the relevance MERLIN+P+TFA prioritization to the system of interest. For each of the remaining regulators (58 of 205 regulators), we tested its association with pluripotency through siRNA knockdown experiments. Many of them were found to have significant positive or negative effect on the pluripotency state of cells (measured by the protein expression of “Nanog” in a cell), thereby establishing them as novel activators or inhibitors of pluripotency. Finally, we experimentally validated the high-confidence target genes of 4 known and 4 novel regulators, using regulator knock down followed by fold change of expression of the targets and demonstrated the importance of using both mRNA and TFA levels of regulators for obtaining high confidence targets.

Our study can be expanded in multiple future directions. Currently TFA estimation is done upstream of GRN inference and the inferred networks are not used to update the inferred TFAs. A direction of future work is to iterate between GRN inference and TFA estimation to expand hidden activity levels to TFs and other regulators without sequence motif information (e.g. TFs with unknown specificity or other regulators, like signaling proteins). In a similar vein, one could incorporate other types of networks, e.g. protein-protein interactions, to expand the prior graph structure to signaling proteins. Another direction of future work is to increase the scalability and non-linear modeling capabilities of MERLIN+P+TFA by incorporating matrix factorization approaches [35, 36]. A third direction of future work is to incorporate other modalities, such as protein levels, for TFA and GRN estimation. The availability of single cell proteomic [37] and multi-modal measurements [38] opens up new opportunities to extend MERLIN+P+TFA to capture these modalities, which could be beneficial for obtaining more causal rather than correlational interactions. In conclusion, we have presented MERLIN+P+TFA that estimates regularized TFA for GRN inference leading to more accurate and comprehensive GRNs across a variety of model systems and for both bulk and single cell RNA-seq datasets. MERLIN+P+TFA and its associated workflow provides a powerful framework for systematic model-based prioritization, validation and discovery of causal regulatory networks in mammalian systems.

## Materials and methods

### MERLIN+P+TFA framework for TFA estimation and gene regulatory network inference

MERLIN+P+TFA combines MERLIN+P’s GRN inference approach [7, 21] with regularized Transcription Factor Activity (TFA) estimation to allow a regulator to be included based on its mRNA or TFA level. Below we describe the original Network Components Analysis (NCA, [14]) approach for TFA estimation, followed by our NCA-LASSO approach regularized version of NCA for more accurate TFA estimation, and finally, the GRN inference procedure of MERLIN+P+TFA.

### Network Component Analysis for TFA estimation

Network component analysis (NCA, [14]) is a method for estimating the unobserved or hidden activity values of TFs given (a) an input graph depicted by an *n* × *m* weighted adjacency matrix, *A*, where each entry *A*(*i, j*) specifies the strength of the connection between gene *i* and TF *j*, and (b) the expression levels of the target genes across a set of experiments, depicted by an *n* × *s* matrix *E*, where each entry *E*(*i, j*) specifies the expression of gene *i* in experiment *j*. *A*(*i, j*) = 0 if *i* is not a target of *j*, and positive otherwise. NCA assumes that the input network structure constrains the TF action on the targets and TFA value can be inferred back by using an alternating least squares optimization approach. Given the expression matrix *E*, NCA decomposes the expression matrix as *E* = *ÂP*, where *P* is the *m* × *s* TFA matrix for *m* TFs across *s* samples. Each entry *P* (*j, l*) specifies the estimated activity of TF *j* in experiment *l*. *Â* is the updated adjacency matrix, having the additional constraint that *Â* should match the sparsity pattern of the input network, *A*. This can be written as the following minimization problem:

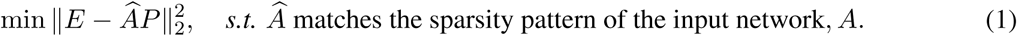

NCA solves this minimization problem by iteratively performing a two-step least square regression. We start by randomly initializing *A*^(0)^ from *A* preserving the zero-pattern for *k* = 0 iteration, and then at each iteration *k >* 0, it first estimates *P* ^(^*^k^*^)^ by keeping *A*^(^*^k^*^−1)^ fixed:

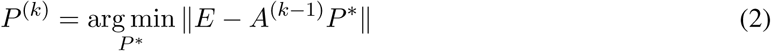

The first step can be solved in closed-form using a pseudoinverse formulation *P* = (*AA*^⊺^)^−1^*A*^⊺^*E*. Next, NCA estimates *A*^(^*^k^*^)^ by keeping *P* ^(^*^k^*^)^ fixed:

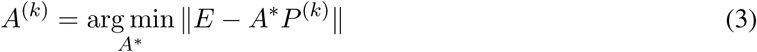

The estimation of the second step, the connectivity matrix *A*^(^*^k^*^)^, can be written as a per-gene regression problem. The *i^th^* row of matrix *A* corresponds to the regression coefficients of all the regulators for target gene *i*. These regression coefficients are estimated for non-zero entries in the original input network, by minimizing the following mean-squared loss:

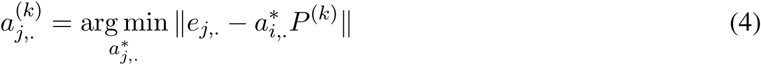

### Regularized TFA estimation

To estimate more robust TFAs from a noisy input network, we incorporate regularization within the NCA framework, which is informed by the weights of TF-target edges in the original input network. It is assumed that higher the weight the more confident is the prior support for the TF-target edge. The original NCA algorithm assumes the input graph is binary and as such ignores these edge weights. In order to incorporate our prior knowledge, we use L1-regularized regression scheme. Let *W* = (*w_ij_*)*_n_*_×_*_m_* represent the penalties on the edges of the input network (the higher the confidence, the lower the penalty). The L1-regularized regression objective we use is:

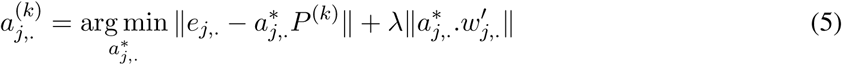

where *λ* is the regularization hyper-parameter. This objective is solved using the “Cyclic Coordinate Descent” algorithm [39]. In the subsequent iterations, this updated *A* matrix is used to update *P* as described above (**Eqn 2**). We run NCA multiple times with different random initializations of *A*^(0)^ to obtain multiple *P* matrices and take the average of these matrices as the final estimated TFA matrix.

### MERLIN+P+TFA network inference

MERLIN+P+TFA infers a genome-scale GRN by combining regularized TFA estimation with MERLIN+P. This is important because the TFA estimation step only prunes the input network and does not add new edges. MERLIN+P+TFA first applies the regularized NCA to estimate the TFA given a prior network and an expression dataset. Then MERLIN+P is employed to infer the regulatory network given the TFA, prior network, and expression dataset. More specifically, let *X_nca_* denote the estimated TFA level for regulator *X*. We appended these estimated TFA profiles for all regulators for which this was estimated allowing a regulator to participate in the GRN based on its mRNA or estimated TFA level, if available. Thus, if regulator “X” was present in both the expression dataset and the estimated TFA profile, it was assigned two rows in the combined (expression+TFA) matrix, one with “X” (its observed expression levels) and another with “X nca” (its estimated TF activities).

MERLIN+P+TFA uses a similar stability selection framework as MERLIN+P which can improve network inference results [40]. Briefly, we create *n* random subsamples of the combined matrix without replacement where each subsample consists of half the number of samples in the combined matrix (*n* = 10 for our simulation studies and 100 for the real data experiments). Subsequently, one regulatory network is inferred from each subsample (with the default setting of MERLIN+P: clustering threshold `-h` = 0.6, sparsity hyper-parameter `-s` = −5, modularity prior `-r` = 4, and prior weight `-q` = 5. Finally, the union of the inferred networks is taken to create a consensus network and each edge is assigned a confidence score which is defined by the proportion of the *n* inferred networks where that edge is present. In addition to inferring the regulatory network, MERLIN+P also reports a co-expression/co-regulation cluster assignment or modules for all target genes [21]. To create a consensus clustering or module assignment, we create a co-clustering similarity matrix (where the (*i, j*)*^th^* element indicates how many times genes *i* and *j* were in the same cluster among the *n* individual runs of MERLIN+P), and apply hierarchical clustering using our in-house programs (average linkage and similarity threshold of 0.3). In our downstream analysis, we only considered the modules with 5 or more members.

### Simulation experiments

The goal of our simulation experiments was two-fold: (1) to examine how noise in the prior network affects the TFA estimation and GRN inference, and (2) to assess improvement in both these tasks when using regularized regression to account for the quality of the prior network. To generate the prior network for the simulation, we randomly selected 100 TFs and 1, 000 genes from the targets of these selected TFs from the yeast motif prior network (**Section Expression datasets**). Subsequently, NCA+LASSO was applied to the prior network and yeast NatVar expression data to estimate the TFA profile (*P*) and regression coefficients for the edges of the prior network (*A*). We generated the simulated expression data (*E*) using *E* = *AP* .

#### Adding noise to the prior network

We simulated different levels of noise in the prior network by removing *X*% of true edges and adding *X*% of false edges at random, where *X*% ∈ {10%, 30%, 50%}, relative to the size of input network (i.e. for a network with 1, 000 edges, 10% noise means removing 100 edges that are present in the network, and adding 100 edges that are not present in the network). For each noise level, we repeated the procedure of adding noise 5 times producing five noisy prior networks for each noise level.

#### Incorporating edge weights in the prior network

We used the beta distribution with parameters *a* and *b* to model edge weights of the simulated prior network, where *a* controls the edge presence probability and *b* controls the edge absence probability (**Figure S1**). We set *a* and *b* to different settings to generate edge weights following two weighting schemes: (1) *Uniform:* All edge weights (for both true and false edges) were set uniformly using samples from a single beta distribution with parameters (*a* = 1, *b* = 1) (**Figure S1A**), (2) *Non-uniform:* We used two different beta distributions for the true and the false edges (**Figure S1B**). For the true edges, (*a* = 5, *b* = 1), enabling the edge presence to be higher probability. For the false edges, we used (*a* = 1, *b* = 5), enabling edge absence to have higher probabilities. If regularization in NCA is helpful, we should observe improvement when using non-uniform edge weights from the second distribution. For each of the 5 noisy prior networks at each noise level, we created two variants of edge-weight distributions that have the same structure but different edge weights. These noisy and edge-weighted prior networks were used for both estimating TFAs and inferring networks.

#### Estimating TFAs and inferring networks

We created 10 subsamples of the simulated expression data, each subsample containing half the samples in each, without replacement. For each subsample and prior network, we first estimated a TFA profile and then inferred a network using the MERLIN+P method (with its default setting). To estimate regularized TFA, we used *λ* ∈ {0, 0.001, 0.005, 0.010, 0.015, 0.020, 0.05, 0.10, 0.15, 0.20}. Here, *λ* = 0 corresponds to the unregularized TFA from the original NCA algorithm.

#### Comparing the estimated TFA to true TFA

For each run of MERLIN+P+TFA on each subsample, we concatenated the estimated TFA profiles of all 100 TFs into a long vector and calculated its absolute Pearson correlation with the true TFA profile. When comparing the performance of one noise scenario to another given a specific edge weighting variant, we used the distribution and median of these correlation values (50 correlation values from 5 replicates of the prior network and 10 expression subsamples, for each noise scenario).

#### Evaluating the inferred networks

We created a consensus network from the 10 networks inferred by MERLIN+P+TFA, where each edge was assigned a confidence value based on the number of times it appeared in the inferred networks. Edges were ranked by their confidence value and given as input to the AUCCalculator Java package [41] along with the gold standard network to calculate the area under the precision-recall curve (AUPR). For each noise level, we estimated 5 AUPR values corresponding to the 5 replicates of the prior network.

### MERLIN+P+TFA for GRN inference from bulk gene expression data

#### Expression datasets

We obtained expression datasets for yeast and mammalian cell lines (mouse and human). The datasets and experimental designs are described below. **Table 2** summarizes the number of samples and datasets for each cell line.

To create the expression matrix of yeast, three natural variation expression datasets (NatVar) were combined from Brem et al. [42], Smith et al. [43], and Zhu et al. [44], as described in Siahpirani et al. [7].

To create the mouse embryonic stem cell (mESC) expression matrix, we used publicly available datasets from the gene expression omnibus (GEO, [45]) database. We searched GEO for expression datasets (both microarray and RNA-seq) measuring expression in mESC or induced pluripotent stem cells (iPSCs). Consequently, we collected 3, 504 microarray samples from 194 datasets, and 2, 182 RNA-seq samples from 216 datasets. For the microarrays, when the raw data was available, the expression was extracted using appropriate R [46] packages depending on the type of the array. On the other hand, when the raw data was not available for the microarrays, we used the processed expression values uploaded to GEO. The RNA-seq data were processed using RSEM [47].

For human cell lines, hESC, MCF7 and LCL, we used MetaSRA [48], which can search the meta data of the sequence read archive (SRA) [49] database for human samples, and GEO for relevant samples. We downloaded 1, 198 samples from 130 datasets for the MCF7 cell line, 1, 501 samples from 43 datasets for the LCL cell line, and 2, 195 samples from 131 datasets for the human ESC (hESC)/iPSC cell line. Since all of these samples used RNA-seq, they were processed with RSEM.

For each of the aforementioned datasets (yeast and mammalian), we collapsed the replicates to mean, and applied quantile normalization (with `quantilenorm()` in MATLAB [50]) to the merged datasets. To remove batch effects, each gene’s expression was zero-meaned per dataset. Moreover, in each dataset, we retained a gene only if its expression is at least (2 × standard deviation) higher than the mean in no less than 6% of the samples.

#### Prior networks

The motif prior network in yeast was obtained from Siahpirani et al. [7]. Briefly, we used position-weight matrices from Gordân et al. [51] and YeTFaSCo [52], and scanned yeast promoters (defined as 1, 000 bp upstream of the first ATG of a gene) using the TestMotif program [53]. The edges were weighted by percentile ranking (defined by motif *p*-value, with highest ranks assigned to lowest *p*-values). To create the prior network for mESC, we used motif instances from the CIS-BP database [54] and the DNase I data of mouse ESC from the ENCODE database [55]. Subsequently, PIQ [56] was applied to find motif instances and score them by their DNase footprint. We mapped motif instances to ±10*Kbp* of the transcription start site (TSS), and selected the motif instances with purity score ≥ 0.85. Finally, the edges were weighted by their percentile ranks based on their motif p-values. For the human cell lines, we used corresponding DNase I data from ENCODE, and in addition to CIS-BP motifs, we also used ENCODE and JASPAR motif instances. For the human cell lines, we used DNase I data from ENCODE and the CIS-BP motif instances as well as the ENCODE [57] and JASPAR [58] motif instances. **Table 3** describes the details of these prior networks.

#### Gold standard networks

For the yeast experiments, we used 4 different gold standard networks: one based on ChIP-chip experiments from MacIsaac et al. [59], two from the YEASTRACT database [60] (*type*≥*2* consists of edges found in two or more different types of experimental assays, and *count* ≥*3*, consists of edges that were detected in three or more experiments), and one KO based networks from Hu et al. [61] (**Table 4**).

For mESC, we used ChIP-based and KD (knock down) based gold standard networks from the ESCAPE database [62] (**File S11**). Moreover, we retrieved the significant knockdown interactions identified in [63–65] and added them to the KD network (**Table 5, File S12**). As another gold standard, we formed a network with the edges that are common between the ChIP and KD gold standard networks.

For hESC, we transformed mESC gold standard networks, ChIP and KD, to hESC gold standard networks ChIP and KD, respectively, by converting the mouse gene names to human gene names. Additionally, we used the ChIP-seq peaks available in ENCODE for the ESC cell line. For each TF, the peaks were mapped to ±10Kbp of TSS of human genes and the resultant TF-target interactions were added to the hESC ChIP gold standard network. For LCL, we used the ChIP and KD interactions reported in Cusanovich et al. [66], and, as an additional gold standard, we used the intersection of the ChIP and KD networks. For MCF7, we obtained the corresponding ChIP-seq peaks from ENCODE and processed them using the same pipeline as that of hESC.

For each gold standard, we only used the edges where both regulator and target have expression in the given dataset (and if a regulator only had TFA we did not use that edge). **Table 6** summarizes the details of these networks.

#### TFA estimation and network inference

For each expression dataset and the corresponding prior network, we used NCA+LASSO to estimate the TFA from all the samples. For the yeast experiments, we used 10 random initializations of *A*^(0)^ (i.e. we estimated TFA 10 times and took the average) along with *λ* ∈ {0.000, 0.005, 0.020, 0.100} for regularization (where *λ* = 0 corresponds to no regularization). For the mammalian experiments, we used 100 random initializations of *A*^(0)^ along with *λ* ∈ {0.000, 0.005, 0.010, 0.020, 0.100} for regularization. For the network inference in yeast, we employed MERLIN+P [7], MERLIN [21] (MERLIN+P without prior), GENIE3 [17], Inferelator [6,18], TIGRESS [19], NetREX [20], and mLASSO-StARS [13]. For the mammalian network inference, only MERLIN and MERLIN+P were employed. For GENIE3, we used *K* = `sqrt` and `nb trees` = 1000. For Inferelator, modified elastic net (MEN) was used with 20 bootstraps where RNA degradation rate (*τ*) was set to 20.1 and prior weight was set to 0.01 (as done in Siahpirani et al. [7]). MERLIN and MERLIN+P were run as described in **Section MERLIN+P+TFA framework for TFA estimation and gene regulatory network inference**. For all inference methods, we used both expression and TFA of regulators when available.

#### Additional evaluation metrics for real expression datasets

##### AUPR, F-score and predictable TFs

To evaluate the inferred networks, we used three evaluation metrics: 1) area under the precision-recall curve (AUPR), 2) F-score and 3) the number of predictable TFs. We used the complete network for AUPR calculation whereas only the top 30*K* of inferred edges were used for F-score computation and counting the number of predictable TFs for the yeast and mammal bulk datasets. Our previous experiments have shown that changing the number of edges (from 10*K* to 50*K*) does not significantly change the results [7]. For the scRNAseq datasets, the top 5*K* of inferred edges were used for F-score computation and counting the number of predictable TFs to be consistent with McCalla et al. [29]. We calculated each AUPR score by comparing an inferred network to a given gold standard network as described in the previous sections. We calculated the F-score for each inferred network which as the harmonic mean of precision and recall obtained from a selected number of top edges in comparison to the gold standard network.

##### Regulator ranking

To assess the importance of a regulator in an inferred network, we determined how much the predictive power of the network decreases in predicting the expression levels of the genes if we remove that regulator from the network, as described in Siahpirani et al. [7] (see **File S13** for inferred network). Briefly, for each target gene, a linear regression model was trained (with 5-fold cross validation) to predict the expression levels of the target gene as a function of that of its regulators (**Figure 7**, **S7**). Subsequently, for each of the regulators of that gene, we removed the regulator and calculated the change in the prediction error. The importance of a regulator was then calculated as the sum of the changes in the prediction errors for all of its targets. To create the list of regulators associated with the ESC state, we used the genes assigned to GO terms “regulation of embryonic development” and “embryonic organ development” in addition to the genes reported by Müller et al. [67] and Wong et al. [68] on MSigDB [69, 70](**Table 7**).

### Experimental validation of MERLIN+P+TFA mESC GRN

#### mESC cell culture

E14 feeder independent mESC and 2D4 feeder dependent iPSC cell lines were maintained in mouse ESC media containing serum.

#### siRNA transfection

Cell lines were plated at a density of 100,000 cells per well in 12-well plates. Reverse transfection was performed with 20 nM of siRNA (purchased from IDT) using DharmaFECT-1 according to the manufacturer’s protocol. Each transcription factor was targeted with a 4 siRNApool (see **File S14**) in quadruplicate. Two rounds of forward transfection were performed at 36 hours with 25 nM siRNA and 60 hours with 40 nM of siRNA to accommodate increased cell numbers from cell division. siRNAs targeting Pou5f1 and Nanog were used as positive controls for pluripotency depletion and non-targeting (NT) siRNA sequence (UAGCGACUAAACACAUCAA) and mock transfected without any siRNA were used as negative controls. Cells were harvested 72 hours after initial transfection to assess knockdown efficiency.

#### RNA extraction, cDNA synthesis, and RT-qPCR

Total RNA was isolated using the ISOLATE II RNA Mini Kit following the manufacturer’s protocol. For each of the tested TFs, including positive and negative controls, three technical replicates were obtained. 1 ug of RNA was converted to cDNA using the qScript cDNA Synthesis Kit. Real time quantitative PCR (qPCR) was for Pou5f1 and Nanog was performed to assess pluripotency with Gapdh and RNA Polymerase II serving as housekeeping controls (see **File S15** for primer details).

#### Flow cytometry

To detect pluripotency protein expression, cells permeabilized using the BD Cytofix/Cytoperm Kit (BD Biosciences) according to manufacturer instructions. Briefly, cells were incubated in BD Cytofix/Cytoperm Reagent for 15-30 minutes at room temperature, washed with BD Perm/Wash buffer, and further treated with

BD Permeabilization Plus buffer on ice for 10 minutes. After an additional wash, intracellular staining was performed using the BD Perm/Wash buffer containing anti-mouse Nanog-eFluor 660 antibodies for 1 hour at room temperature. Cells were washed, pelleted, and resuspended in BD Perm/Wash buffer for analysis. Samples were quantified on a Flow Cytometer and analyzed using FlowJo Software.

#### Target validation of novel TFs

To further validate TFA prediction accuracy and measure the effect of the novel TFs on pluripotency, we selected a subset of the novel TFs tested with siRNA knockdown to assess the effect of the siRNA on the predicted target genes of the novel TFs. Specifically, we investigated four novel TFs not present in the mESC gold standard networks (Fosl1, Gtf2a2, Zbtb7b, Lztr1), along with four of the well-characterized TFs (Esrrb, Nanog, Oct4, Zfp281) serving as positive controls. Cell culture and siRNA transfection were conducted in the E14 cell line with each condition performed in duplicate to obtain a total of three biological replicates for both novel and known TFs. Following RNA extraction and cDNA synthesis steps as described for earlier siRNA sample processing, qPCR was performed in duplicate for the six tested TFs with primers designed to measure expression of targets for the TF undergoing siRNA knockdown (see **File S15** for primer details).

### MERLIN+P+TFA for GRN inference from scRNAseq datasets

We applied MERLIN+P+TFA, Inferelator with prior (Inferelator+P) and SCENIC network inference algorithms on scRNA-seq datasets from our previous benchmarking study of scRNA-seq GRN inference algorithms by McCalla et al. [29]. We selected these three algorithms for further investigation as MERLIN and Inferelator can utilize prior network inputs, while SCENIC was highly ranked with reasonable computational runtimes in a previous scRNAseq network inference benchmarking study [29]. For regularized TFA, we used *λ* ∈ {0.005, 0.010, 0.020, 0.100}. We evaluated the performance of regularized TFA for scRNAseq datasets when applied with the MERLIN [21] [7], Inferelator [30], and SCENIC [31] network inference algorithms. We utilized the previously inferred scRNAseq networks from McCalla et al. [29] that compared networks inferred from expression alone, the addition of priors, TFA generated using NCA (unregularized TFA), and Inferelator’s in-algorithm estimated TFA.

The datasets we considered were from three different organisms: human dendritic cells (Shalek [71]), human pluripotent stem cells (Han), yeast (Gasch [32] and Jackson [30]), and two mouse reprogramming datasets (Sridharan FBS [72], Sridharan A2S [72]). Preprocessed datasets and input priors were prepared as detailed in McCalla et. al [29]. The inferred GRNs were evaluated using F-score, AUPR, and predictable TFs similar to our bulk RNA-seq experiments using the gold standard networks Perturb, ChIP, and the intersection of Perturb and ChIP as described in McCalla et al. [29]. For the output of each algorithm that utilized regularized TFA, we selected the one best inferred network out of the four regularized TFA networks that returned the best AUPR result when testing regularized TFA settings of *λ* ∈ {0.005, 0.010, 0.020, 0.100}.

This “best regularized TFA” network was used for network metric calculations for both AUPR and predictable TFs.

## Supporting information

Manuscript tables

Supplementary figures, tables, and file list

## Acknowledgments

We thank the Center for High Throughput Computing (CHTC) at UW-Madison for computational resources. We thank Khoa Tran for providing the list of ESC associated genes and help in reviewing mouse ESC samples. We thank the members of the Roy and Sridharan groups for helpful discussions. This work was supported by NSF grants IOS 1840687 and IOS 2406533, NIH grants R01GM144708 and R01GM117339, and DOE grant DE-SC0021052 to S.R., R01HD105151 to R.S. and a UW-Madison Stem Cell and Regenerative Medicine Center predoctoral fellowship to C.M.D.

