## Supplementary material for "Uncovering Functional Gene Regulatory Networks in Bulk and Single-Cell Data through Robust Transcription Factor Activity Estimation and Model-Guided Experimental Validation": Manuscript tables

| | 2D4 Cell line | | E14 Cell line | |
|---|---|---|---|---|
| | NT<TF | NT>TF | NT<TF | NT>TF |
| Zbtb7b | 4.65E-62 | 0.239 | 3.08E-212 | 0.0055 |
| Gtf2a2 | 3.72E-32 | 0.934 | 4.76E-152 | 0.997 |
| Ehf | 2.34E-82 | 0.998 | 1.49E-93 | 1 |
| Foxc1 | 2.25E-71 | 0.983 | 1.17E-90 | 0.212 |
| Dmrta2 | 1.63E-149 | 0.995 | 7.32E-54 | 1 |
| Fosl1 | 2.91E-06 | 1.94E-21 | 3.01E-38 | 1.47E-13 |
| Creb3 | 0.896 | 2.12E-06 | 1.04E-33 | 1 |
| Asz1 | 5.93E-119 | 0.99 | 5.11E-32 | 0.985 |
| Mypop | 3.56E-25 | 0.981 | 8.98E-29 | 0.156 |
| Zfp281 | 5.7043e-20 | 5.1030e-02 | 2.0139e-28 | 9.9909e-01 |
| Gmeb1 | 3.95E-09 | 0.0287 | 5.53E-25 | 0.96 |
| Lztr1 | 1.1E-28 | 0.996 | 1.85E-22 | 1 |
| Nelfa | 0.706 | 0.49 | 3.3E-20 | 0.000235 |
| Zfp784 | 0.00323 | 0.74 | 6.97E-19 | 3.03E-15 |
| Zfp110 | 0.996 | 1.35E-17 | 2.24E-17 | 0.00953 |
| Esrrb | 2.25E-296 | 1 | 4.19E-15 | 0.983 |
| Hmbox1 | 4.58E-10 | 0.708 | 8.44E-13 | 2.04E-05 |
| Arx | 1.95E-10 | 1 | 3.85E-12 | 0.996 |
| Hsfy2 | 6.73E-19 | 0.369 | 1.53E-09 | 1 |
| Xbp1 | 0.886 | 1.05E-21 | 2.5E-09 | 5.91E-22 |
| Dux | 0.0822 | 0.0765 | 3.73E-09 | 2.65E-07 |
| Creb3l3 | 0.685 | 4.84E-48 | 3.77E-07 | 0.269 |
| Batf3 | 3.66E-47 | 0.946 | 1.51E-05 | 0.998 |
| Mecp2 | 0.00543 | 1.55E-05 | 1.57E-05 | 0.935 |
| Churc1 | 0.862 | 0.00808 | 0.000223 | 0.961 |

| | | | | |
|---|---|---|---|---|
| Bhlha15 | 0.000191 | 1 | 0.000447 | 0.965 |
| Gmeb2 | 2.49E-17 | 0.787 | 0.00128 | 0.857 |
| Dtx3l | 0.978 | 2.4E-14 | 0.00202 | 0.458 |
| Elf1 | 0.893 | 4.77E-17 | 0.00252 | 0.0288 |
| E2f6 | 9.06E-10 | 0.919 | 0.0307 | 3.77E-27 |
| Six4 | 0.347 | 0.107 | 0.049 | 1.78E-06 |
| Zbtb7a | - | - | 0.0744 | 1.54E-24 |
| Hey1 | 7.55E-16 | 1 | 0.123 | 3.63E-16 |
| Lin54 | 4.87E-104 | 4.48E-38 | 0.142 | 2.71E-128 |
| Noct | 0.105 | 0.000213 | 0.176 | 0.453 |
| Creb3l2 | 0.141 | 3.41E-19 | 0.591 | 1.07E-20 |
| Rhox6 | 5.74E-05 | 0.968 | 0.627 | 1.68E-40 |
| Foxc2 | 0.87 | 6.38E-16 | 0.642 | 9.82E-33 |
| Scml2 | 0.00447 | 0.743 | 0.753 | 3.8E-82 |
| Kdm5d | 1.29E-20 | 0.946 | 0.834 | 9.54E-30 |
| Gcm2 | 0.158 | 0.0239 | 0.853 | 6.27E-29 |
| Hes7 | 0.0657 | 0.018 | 0.865 | 0.0432 |
| Enpp3 | 3.56E-10 | 0.917 | 0.883 | 1.08E-29 |
| Sp3 | 0.0679 | 0.0605 | 0.914 | 4.77E-10 |
| Foxj3 | 0.996 | 2.74E-07 | 0.933 | 1.64E-55 |
| Col5a2 | 0.88 | 2.11E-18 | 0.946 | 3.28E-95 |
| Scrt2 | 0.916 | 3.91E-25 | 0.971 | 3.32E-41 |
| Mlxip | 0.673 | 5.09E-08 | 0.974 | 1.58E-169 |
| Foxk2 | 2.39E-23 | 0.811 | 0.987 | 2.18E-14 |
| Bhlhe40 | 0.0285 | 0.0549 | 0.993 | 0.000702 |
| Scrt1 | 0.902 | 0.129 | 0.998 | 2.51E-50 |
| Tcfl5 | 0.123 | 0.313 | 0.999 | 8.21E-70 |

| | | | | |
|---|---|---|---|---|
| Bach1 | 0.955 | 1.72E-09 | 1 | 1.07E-08 |
| Atf1 | 0.925 | 0.00058 | 1 | 4.57E-90 |
| Tbx20 | 1.67E-08 | 0.985 | 1 | 2.22E-160 |
| Ankrd22 | 0.977 | 4.92E-17 | - | - |
| Alx4 | 2.41E-11 | 0.0388 | - | - |
| Aes | 2.13E-35 | 0.000254 | - | - |

**Table 1.** The $p$-values of two-sample Kolmogorov–Smirnov (KS) test. There is a pair of $p$-values referred to as "NT<TF" and "NT>TF" for each TF (such as Zbtb7b) with respect to each cell line (2D4 or E14). The "NT<TF" $p$-value indicates whether the cumulative distribution function (CDF) of the fluorescent measurements after the knockdown of the TF in that cell line is significantly larger than the CDF of the fluorescent measurements of the "non-targeting siRNA control" (NT) in that cell line. If the $p$-value is significant (i.e. less than $1e-10$), then it is colored in red (Zbtb7b in the 2D4 cell line for instance), indicating that we can reject the null hypothesis that the knockdown CDF and control CDF are coming from the same continuous distribution in favor of the alternative hypothesis that the knockdown CDF is coming from a larger continuous distribution than that of the control CDF. In contrast, the "NT<TF" $p$-value indicates whether the knockdown CDF is smaller than the control CDF or not. If the $p$-value is significant (i.e. less than $1e-10$), then it is colored in green (Fosl1 in the 2D4 cell line for instance). The corresponding histograms and cumulative distribution function (CDF) plots are presented in supplementary file "sfile6_KS_test_p_vals.xlsx."

| | #datasets | #samples (before collapsing replicates) | #samples (after collapsing replicates) | #genes | #regs with Exp | #regs with TFA | #regs with both |
|---|---|---|---|---|---|---|---|
| Yeast (NatVar) | 3 | 621 | 377 | 5,660 | 537 | 197 | 197 |
| mESC/ iPSC (Array) | 171 | 3,504 | 1,178 | 4,561 | 917 | 252 | 113 |
| mESC/ iPSC (RNA-seq) | 216 | 2,182 | 764 | 4,743 | 926 | 258 | 105 |
| MCF7 (RNA-seq) | 130 | 1,198 | 534 | 5,513 | 635 | 420 | 72 |
| Human LCL (RNA-seq) | 43 | 1,501 | 1,000 | 5,614 | 833 | 447 | 184 |
| hESC/ iPSC (RNA-seq) | 131 | 2,195 | 1,316 | 5,756 | 998 | 420 | 171 |

**Table 2.** Statistics on yeast and mammalian expression datasets. Number of datasets, samples (before and after collapsing replicates), number of genes, number of regulators with expression, with estimated TFA, or both expression and TFA, used for each cell line.

| | Exp edges | | | TFA edges | | |
|---|---|---|---|---|---|---|
| | #Regs | #Targets | #Edges | #Regs | #Targets | #Edges |
| Yeast (NatVar) | 197 | 5,506 | 187,079 | 197 | 5,506 | 187,079 |
| mESC/iPSC (Array) | 331 | 4,496 | 329,763 | 252 | 4,508 | 242,735 |
| mESC/iPSC (RNA-seq) | 278 | 4,591 | 313,184 | 258 | 4,615 | 282,076 |
| MCF7 (RNA-seq) | 144 | 5,377 | 275,512 | 420 | 5,432 | 861,664 |
| Human LCL (RNA-seq) | 279 | 5,508 | 653,316 | 447 | 5,516 | 927,195 |
| hESC/iPSC (RNA-seq) | 310 | 5,684 | 600,814 | 420 | 5,693 | 850,268 |

**Table 3.** Number of regulators, targets, and edges in each of the prior networks, for expression-regulator (Exp) edges (edges where we have the expression levels of the regulator) or TFA-regulator (TFA) edges (edges where we have the estimated TFA of the regulator).

| Gold standard | All Regs | All PTFs | Exp-based | Prior-based | Exp- & Prior-based | Exp-based + TFA | Prior-based + TFA | Exp- & Prior-based + TFA |
|---|---|---|---|---|---|---|---|---|
| MacIsaac | 116 | 61 | 20 | 50 | 50 | 37 | 57 | 59 |
| YEASTRACT (*type*$\geq$*2*) | 127 | 68 | 31 | 56 | 58 | 51 | 62 | 65 |
| YEASTRACT (*count* $\geq$*3*) | 120 | 68 | 32 | 57 | 60 | 48 | 63 | 65 |
| Hu KO | 268 | 37 | 12 | 12 | 19 | 19 | 26 | 30 |

**Table 4.** Number of regulators, and predictable TFs for gold standard networks in yeast. Each row corresponds to a specific gold standard network and each column corresponds to a specific group of inferred networks. The only exception is column "All Regs" which simply presents the total number of regulators in the corresponding gold standard network. Column "All PTFs" represents the total number of predictable TFs across all the inferred networks and the corresponding prior network. Here, the inferred networks refer to the networks inferred with methods {GENIE3, Inferelator, MERLIN, MERLIN+P} for five different TFA settings–no TFA i.e. using expression data only (Exp), TFA estimated with NCA (TFA0.000), TFA estimated with regularized NCA having $\lambda = 0.005$ (TFA0.005), $\lambda = 0.02$ (TFA0.020), $\lambda = 0.1$ (TFA0.100). Column "Exp-based" represents the number of predictable TFs in the networks inferred with the expression-based methods (i.e. the methods that can not explicitly incorporate prior network during network inference, namely, {GENIE3, MERLIN}) for the Exp setting. Column "Prior-based" represents the number of predictable TFs in the networks inferred with the prior-based methods (i.e. the methods that can explicitly incorporate prior network during network inference, namely, {Inferelator, MERLIN+P}) for the Exp setting. Column "Exp- & Prior-based" represents the number of predictable TFs in the networks inferred with both expression-based and prior-based methods for the Exp setting. Column "Exp-based + TFA" represents the number of predictable TFs in the networks inferred with the expression-based methods for the {TFA0.000, TFA0.005, TFA0.020, TFA0.100} settings. Column "Prior-based + TFA" represents the number of predictable TFs in the networks inferred with the prior-based methods for the {TFA0.000, TFA0.005, TFA0.020, TFA0.100} settings. Column "Exp- & Prior-based + TFA" represents the number of predictable TFs in the networks inferred with both expression-based and prior-based methods for the {TFA0.000, TFA0.005, TFA0.020, TFA0.100} settings. The top 30,000 edges by edge confidence were considered for counting the predictable TFs in each of the networks.

| Dataset | Gold standard | All Regs | All PTFs | Exp-based | Prior-based | Exp- & Prior-based | Exp-based + TFA | Prior-based + TFA | Exp- & Prior-based + TFA |
|---|---|---|---|---|---|---|---|---|---|
| mESC (RNA-seq) | ChIP | 43 | 30 | 19 | 26 | 28 | 24 | 27 | 30 |
| | KD | 106 | 39 | 19 | 9 | 23 | 34 | 19 | 35 |
| mESC (array) | ChIP | 42 | 31 | 26 | 27 | 30 | 25 | 28 | 31 |
| | KD | 111 | 39 | 31 | 18 | 33 | 33 | 31 | 38 |

**Table 5.** Number of regulators, and predictable TFs for gold standard networks in the mESC cell line. Each row corresponds to a distinct gold-standard network and each column corresponds to a specific group of inferred networks. The only exception is column "All Regs" which simply presents the total number of regulators in the corresponding gold standard network. Column "All PTFs" represents the total number of predictable TFs across all the inferred networks and the corresponding prior network. Here, the inferred networks refer to the networks inferred with methods {MERLIN, MERLIN+P} for six different TFA settings–no TFA i.e. using expression data only (Exp), TFA estimated with NCA (TFA0.000), TFA estimated with regularized NCA having $\lambda = 0.005$ (TFA0.005), $\lambda = 0.01$ (TFA0.010), $\lambda = 0.02$ (TFA0.020), $\lambda = 0.1$ (TFA0.100). Column "Exp-based" represents the number of predictable TFs in the networks inferred with the expression-based method for the Exp setting. Column "Prior-based" represents the number of predictable TFs in the networks inferred with the prior-based method for the Exp setting. Column "Exp- & Prior-based" represents the number of predictable TFs in the networks inferred with both expression-based and prior-based methods for the Exp setting. Column "Exp-based + TFA" represents the number of predictable TFs in the networks inferred with the expression-based method for the {TFA0.000, TFA0.005, TFA0.010, TFA0.020, TFA0.100} settings. Column "Prior-based + TFA" represents the number of predictable TFs in the networks inferred with the prior-based method for the {TFA0.000, TFA0.005, TFA0.010, TFA0.020, TFA0.100} settings. Column "Exp- & Prior-based + TFA" represents the number of predictable TFs in the networks inferred with both expression-based and prior-based methods for the {TFA0.000, TFA0.005, TFA0.010, TFA0.020, TFA0.100} settings. The top 30,000 edges by edge confidence were considered for counting the predictable TFs in each of the networks.

| Dataset | Gold standard | #Regs | #Targets | #Edges |
|---|---|---|---|---|
| Yeast NatVar | MacIsaac | 114 | 1,883 | 3,802 |
| | YEASTRACT (*type*$\geq$2) | 119 | 2,167 | 4,219 |
| | YEASTRACT (*count* $\geq$3) | 119 | 2,067 | 3,818 |
| | Hu | 264 | 2,334 | 10,101 |
| mESC (array) | ChIP | 42 | 3,791 | 22,335 |
| | Loss of function | 52 | 2,619 | 7,116 |
| | Gain of function | 82 | 3,535 | 36,605 |
| | ChIP&LOF overlap | 23 | 1,228 | 2,295 |
| | ChIP&GOF overlap | 16 | 1,225 | 1,947 |
| mESC (RNA-seq) | ChIP | 43 | 3,225 | 21,086 |
| | Loss of function | 51 | 2,422 | 6,625 |
| | Gain of function | 81 | 2,986 | 30,728 |
| | ChIP&LOF overlap | 23 | 1,108 | 2,113 |
| | ChIP&GOF overlap | 18 | 1,088 | 1,802 |
| hESC | ChIP | 67 | 5,324 | 84,637 |
| | KD | 103 | 2,998 | 29,555 |
| | ChIP&KD overlap | 32 | 1,444 | 3,330 |
| human LCL | ChIP | 180 | 2,254 | 103,081 |
| | KD | 58 | 2,246 | 26,881 |
| | ChIP&KD overlap | 29 | 1,715 | 5,328 |
| MCF7 | ChIP | 16 | 4,897 | 29,192 |

**Table 6.** Number of regulators, targets, and edges in each gold standard network of each dataset.

| Module ID | No. of enriched GO terms | No. of enriched regulators |
|---|---|---|
| Cluster3336 | 239 | 9 |
| Cluster2945 | 201 | 3 |
| Cluster3343 | 117 | 6 |
| Cluster3299 | 85 | 0 |
| Cluster3112 | 67 | 2 |
| Cluster3329 | 52 | 2 |
| Cluster3001 | 40 | 2 |
| Cluster3303 | 37 | 1 |
| Cluster3025 | 34 | 4 |
| Cluster3307 | 23 | 1 |
| Cluster3253 | 22 | 1 |
| Cluster2907 | 20 | 2 |
| Cluster3304 | 20 | 1 |
| Cluster3002 | 19 | 2 |
| Cluster3153 | 16 | 13 |
| Cluster3070 | 15 | 2 |
| Cluster3298 | 12 | 3 |
| Cluster3068 | 11 | 1 |
| Cluster2979 | 9 | 0 |
| Cluster3215 | 9 | 4 |
| Cluster3125 | 7 | 2 |
| Cluster3213 | 5 | 1 |
| Cluster3234 | 3 | 2 |
| Cluster3099 | 2 | 0 |
| Cluster3257 | 2 | 0 |
| Cluster3328 | 2 | 1 |

| | | |
|---|---|---|
| Cluster3154 | 1 | 0 |
| Cluster3232 | 1 | 0 |
| Cluster2885 | 0 | 2 |
| Cluster2890 | 0 | 2 |
| Cluster2935 | 0 | 3 |
| Cluster2951 | 0 | 2 |
| Cluster2957 | 0 | 1 |
| Cluster2971 | 0 | 1 |
| Cluster2976 | 0 | 3 |
| Cluster2985 | 0 | 2 |
| Cluster2996 | 0 | 2 |
| Cluster3007 | 0 | 1 |
| Cluster3039 | 0 | 1 |
| Cluster3040 | 0 | 1 |
| Cluster3082 | 0 | 6 |
| Cluster3124 | 0 | 1 |
| Cluster3146 | 0 | 1 |
| Cluster3189 | 0 | 1 |
| Cluster3194 | 0 | 2 |
| Cluster3211 | 0 | 5 |
| Cluster3233 | 0 | 1 |
| Cluster3251 | 0 | 1 |
| Cluster3252 | 0 | 1 |
| Cluster3259 | 0 | 2 |
| Cluster3260 | 0 | 2 |
| Cluster3274 | 0 | 4 |
| Cluster3275 | 0 | 7 |

| | | |
|---|---|---|
| Cluster3282 | 0 | 1 |
| Cluster3300 | 0 | 3 |
| Cluster3308 | 0 | 1 |
| Cluster3309 | 0 | 5 |
| Cluster3335 | 0 | 1 |
| Cluster3339 | 0 | 1 |
| Cluster3341 | 0 | 2 |

**Table 7.** There are a total of 65 gene modules in the network predicted by MERLIN+P+TFA (TFA0.010) from the mESC RNA-seq dataset. We performed gene ontology (GO) enrichment and regulator enrichment for the modules having five or more genes (there are 60 such modules as presented in this table). Here, we present the number of GO terms and number of regulators enriched for each of the modules. The modules are sorted by the descending of the number of enriched GO terms. Six of the modules have no enriched regulators (colored in red). We generated various heatmaps for the remaining 54 modules which are presented in Supplementary ( "S07_merlinp_rnaseq_tfa0010_hmap_out_0_8_0_3.zip").
