## Supplementary figures, tables, and file list for "Uncovering Functional Gene Regulatory Networks in Bulk and Single-Cell Data through Robust Transcription Factor Activity Estimation and Model-Guided Experimental Validation"

### Supplementary figures, tables, and files

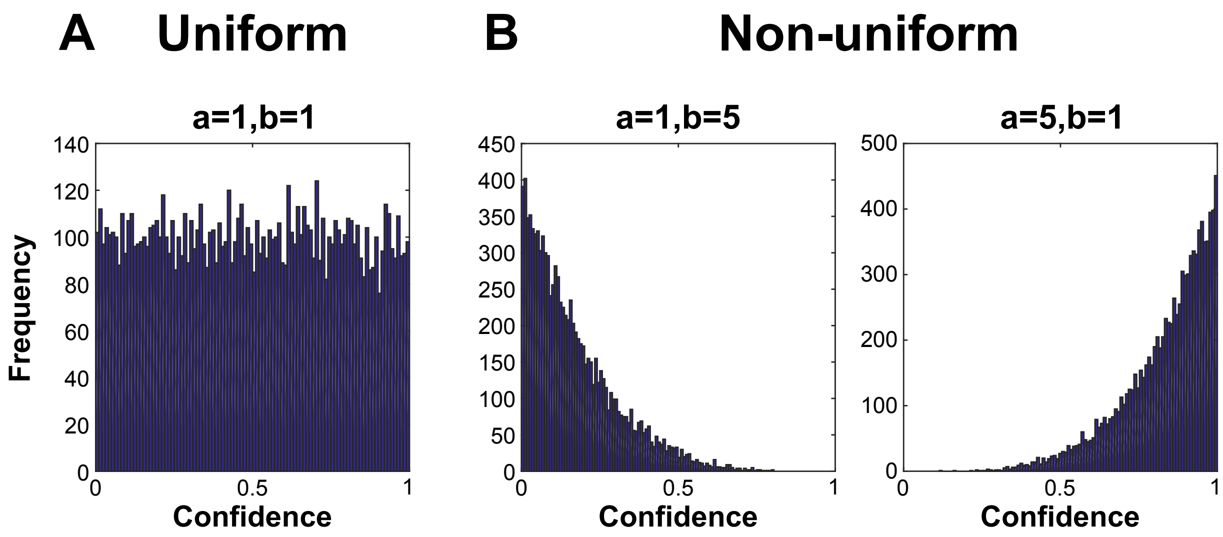
A
Uniform
B
Non-uniform
a=1,b=1
a=1,b=5
a=5,b=1
Frequency
Confidence
Confidence
Confidence
0
20
40
60
80
100
120
140
0
50
100
150
200
250
300
350
400
450
0
100
200
300
400
500
0
0.5
1

**Figure S1.** Distribution of simulated edge weights with different configurations of Beta distribution **A**) $a = 1, b = 1$, uniform. **B**) $a = 1, b = 5$, non-uniform, false edges, and $a = 5, b = 1$, non-uniform, true edges.

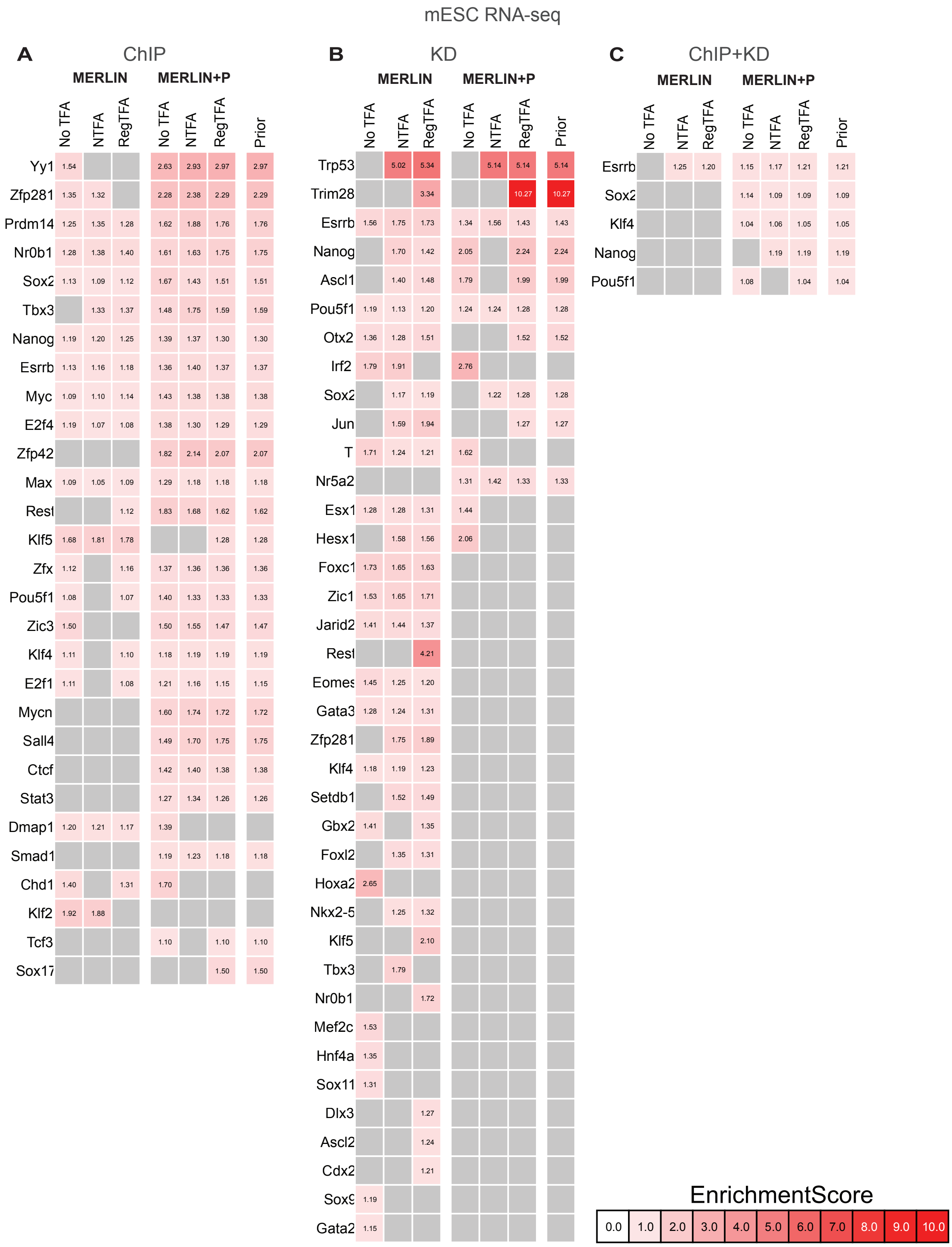
mESC RNA-seq
A
ChIP
MERLIN
MERLIN+P
No TFA
NTFA
RegTFA
Prior
Yy1
Zfp281
Prdm14
Nr0b1
Sox2
Tbx3
Nanog
Esrrb
Myc
E2f4
Zfp42
Max
Rest
Klf5
Zfx
Pou5f1
Zic3
Klf4
E2f1
Mycn
Sall4
Ctcf
Stat3
Dmap1
Smad1
Chd1
Klf2
Tcf3
Sox17
B
KD
Trp53
Trim28
Esrrb
Nanog
Ascl1
Pou5f1
Otx2
Irf2
Sox2
Jun
T
Nr5a2
Esx1
Hesx1
Foxc1
Zic1
Jarid2
Rest
Eomes
Gata3
Zfp281
Klf4
Setdb1
Gbx2
Foxl2
Hoxa2
Nkx2-5
Klf5
Tbx3
Nr0b1
Mef2c
Hnf4a
Sox11
Dlx3
Ascl2
Cdx2
Sox9
Gata2
C
ChIP+KD
Esrrb
Sox2
Klf4
Nanog
Pou5f1
EnrichmentScore
0.0
1.0
2.0
3.0
4.0
5.0
6.0
7.0
8.0
9.0
10.0

**Figure S2.** List of all predictable TFs when recovering **A**) ChIP, **B**) KD, and **C**) the intersection of the ChIP and KD. Each column correspond to one of inferred networks (MERLIN and MERLIN+P) using expression and TFA (at different regularization levels) of regulators. 'RegTFA' represents the best performing regularized NCA result after testing $\lambda = 0.005, 0.010, 0.020, 0.100$. The higher the enrichment score, the more significant the overlap is between the targets of that TF in that particular inferred network and the corresponding gold standard network.

mESC array

**A** ChIP

| | MERLIN | | | MERLIN+P | | | |
|---|---|---|---|---|---|---|---|
| | No TFA | NTFA | RegTFA | No TFA | NTFA | RegTFA | Prior |
| Zfp281 | 1.49 | 1.51 | 1.58 | 2.37 | 2.40 | 2.25 | 2.25 |
| Dmap1 | 1.56 | 1.87 | 1.64 | 1.94 | 1.55 | 1.81 | 1.81 |
| Nr0b1 | 1.41 | 1.63 | 1.72 | 1.80 | 1.84 | 1.82 | 1.82 |
| Mycn | 1.34 | 1.49 | 1.41 | 1.92 | 1.94 | 1.77 | 1.77 |
| Klf5 | 2.47 | 3.10 | 2.95 | 1.50 | 1.41 | | |
| Zfx | 1.63 | 1.56 | 1.56 | 1.59 | 1.63 | 1.55 | 1.55 |
| Zic3 | 1.57 | | 1.99 | 1.67 | 1.75 | 1.71 | 1.71 |
| Tbx3 | 1.42 | 1.46 | 1.51 | 1.48 | 1.38 | 1.56 | 1.56 |
| Rest | 1.13 | 1.17 | 1.18 | 1.92 | 1.67 | 1.60 | 1.60 |
| Myc | 1.36 | 1.32 | 1.31 | 1.63 | 1.51 | 1.52 | 1.52 |
| Yy1 | | | | 2.31 | 2.68 | 2.40 | 2.40 |
| Sox2 | 1.20 | 1.22 | 1.24 | 1.63 | 1.49 | 1.47 | 1.47 |
| Pou5f1 | 1.27 | 1.15 | 1.23 | 1.58 | 1.42 | 1.47 | 1.47 |
| Klf4 | 1.34 | 1.47 | 1.44 | 1.32 | 1.34 | 1.30 | 1.30 |
| E2f4 | 1.25 | 1.10 | 1.20 | 1.47 | 1.38 | 1.41 | 1.41 |
| E2f1 | 1.39 | 1.27 | 1.27 | 1.39 | 1.26 | 1.28 | 1.28 |
| Max | 1.26 | 1.25 | 1.22 | 1.37 | 1.34 | 1.32 | 1.32 |
| Stat3 | 1.22 | 1.15 | 1.20 | 1.47 | 1.39 | 1.30 | 1.30 |
| Esrrb | 1.20 | 1.17 | 1.15 | 1.35 | 1.31 | 1.29 | 1.29 |
| Nanog | 1.16 | 1.17 | 1.18 | 1.22 | 1.23 | 1.25 | 1.25 |
| Zfp42 | | | | 1.79 | 1.85 | 1.75 | 1.75 |
| Trim28 | 1.17 | 1.27 | 1.25 | 1.30 | 1.31 | | |
| Tcf3 | 1.11 | | | 1.15 | 1.14 | 1.10 | 1.10 |
| Ctcf | | | | 1.49 | 1.33 | 1.33 | 1.33 |
| Chd1 | 1.38 | 1.45 | | 1.60 | | | |
| Klf2 | 1.81 | 2.22 | | | | | |
| Smad1 | | | | 1.29 | | 1.21 | 1.21 |
| Setdb1 | 1.11 | | | | | 1.28 | 1.28 |
| Cdx2 | | 1.51 | | | 1.77 | | |
| Sall4 | 1.24 | | | 1.53 | | | |
| Nacc1 | 1.33 | | 1.40 | | | | |

**B** KD

| | MERLIN | | | MERLIN+P | | | |
|---|---|---|---|---|---|---|---|
| | No TFA | NTFA | RegTFA | No TFA | NTFA | RegTFA | Prior |
| Rest | 4.92 | 4.46 | 3.90 | 7.50 | 7.39 | 5.65 | 5.65 |
| Zfp281 | 3.40 | 3.21 | 2.89 | 2.59 | 2.89 | 3.00 | 3.00 |
| Tbx3 | 2.29 | 2.58 | 2.84 | 1.68 | 2.07 | 2.13 | 2.13 |
| Trp53 | | 2.37 | 2.80 | | 2.72 | 3.38 | 3.38 |
| Cbx8 | 1.71 | 2.27 | 2.08 | 2.19 | 3.81 | | |
| Klf5 | 3.12 | 3.58 | 3.68 | | 1.45 | | |
| Nanog | 1.42 | 1.53 | 1.57 | 1.70 | 1.84 | 1.68 | 1.68 |
| Esx1 | 1.28 | 1.34 | 1.47 | 1.53 | 1.62 | 1.58 | 1.58 |
| Esrrb | 1.52 | 1.71 | 1.71 | 1.28 | 1.45 | 1.35 | 1.35 |
| Gata2 | 1.50 | 1.63 | 1.60 | 1.18 | 1.24 | 1.24 | 1.24 |
| Ascl2 | 1.30 | 1.45 | 1.40 | | 1.84 | 1.73 | 1.73 |
| Pou5f1 | 1.41 | 1.24 | 1.42 | 1.40 | 1.29 | 1.35 | 1.35 |
| Sox2 | 1.29 | 1.31 | 1.42 | 1.30 | 1.36 | 1.38 | 1.38 |
| Jun | 1.51 | 1.46 | 1.51 | 1.21 | 1.21 | 1.25 | 1.25 |
| T | 1.58 | 1.22 | 1.40 | 1.67 | | 1.56 | 1.56 |
| Bcl6 | 1.34 | 1.39 | 1.49 | | 1.58 | 1.56 | 1.56 |
| Gbx2 | 1.59 | 1.55 | 1.69 | 1.60 | 1.54 | | |
| Klf4 | 1.43 | 1.46 | 1.50 | | 1.16 | 1.18 | 1.18 |
| Cdx2 | 1.21 | 1.23 | 1.26 | 1.40 | | 1.34 | 1.34 |
| Gata3 | 1.28 | 1.38 | 1.37 | | 1.14 | 1.15 | 1.15 |
| Fosl2 | 1.79 | 2.14 | 1.88 | | | | |
| Nr0b1 | 1.71 | | 1.81 | 2.27 | | | |
| Setdb1 | 1.70 | 1.87 | 2.18 | | | | |
| Foxc1 | 1.30 | 1.55 | 1.49 | | 1.20 | | |
| Msc | 1.25 | | 1.28 | | | 1.45 | 1.45 |
| Gadd45a | | | | | | 2.60 | 2.60 |
| Sox11 | 1.48 | 1.42 | 1.51 | | | | |
| Otx2 | 1.39 | 1.28 | | | 1.34 | | |
| Nr5a2 | | | | 1.30 | | 1.33 | 1.33 |
| Sox9 | 1.16 | | 1.23 | | 1.39 | | |
| Eomes | 1.30 | 1.16 | 1.17 | | | | |
| Mef2c | 1.40 | 1.51 | | | | | |
| Zic1 | 1.62 | | | 1.27 | | | |
| Elf1 | 1.39 | 1.24 | | | | | |
| Hoxa2 | | | | 2.45 | | | |
| Foxl2 | | | | | | 1.12 | 1.12 |
| Jarid2 | | | 1.41 | | | | |

**C** ChIP+KD

| | MERLIN | | | MERLIN+P | | | |
|---|---|---|---|---|---|---|---|
| | No TFA | NTFA | RegTFA | No TFA | NTFA | RegTFA | Prior |
| Klf4 | | 1.13 | 1.10 | 1.08 | 1.07 | 1.09 | 1.09 |
| Sox2 | | 1.06 | 1.06 | 1.13 | 1.06 | 1.07 | 1.07 |
| Pou5f1 | | 1.04 | 1.08 | 1.09 | 1.04 | 1.09 | 1.09 |
| Setdb1 | 1.33 | 1.43 | 1.28 | 1.31 | | | |
| Nanog | 1.11 | | | 1.16 | | 1.14 | 1.14 |
| Tbx3 | | 1.41 | | 1.48 | 1.40 | | |
| Myc | | 1.14 | | 1.22 | 1.17 | | |
| Sox17 | | 1.43 | | | | | |

EnrichmentScore

| | | | | | | | | | | |
|---|---|---|---|---|---|---|---|---|---|---|
| 0.0 | 1.0 | 2.0 | 3.0 | 4.0 | 5.0 | 6.0 | 7.0 | 8.0 | 9.0 | 10.0 |

**Figure S3.** List of all predictable TFs when recovering **A**) ChIP, **B**) KD, and **C**) the intersection of the ChIP and KD. Each column correspond to one of inferred networks (MERLIN and MERLIN+P) using expression and TFA (at different regularization levels) of regulators. 'RegTFA' represents the best performing regularized NCA result after testing $\lambda = 0.005, 0.010, 0.020, 0.100$. The higher the enrichment score, the more significant the overlap is between the targets of that TF in that particular inferred network and the corresponding gold standard network.

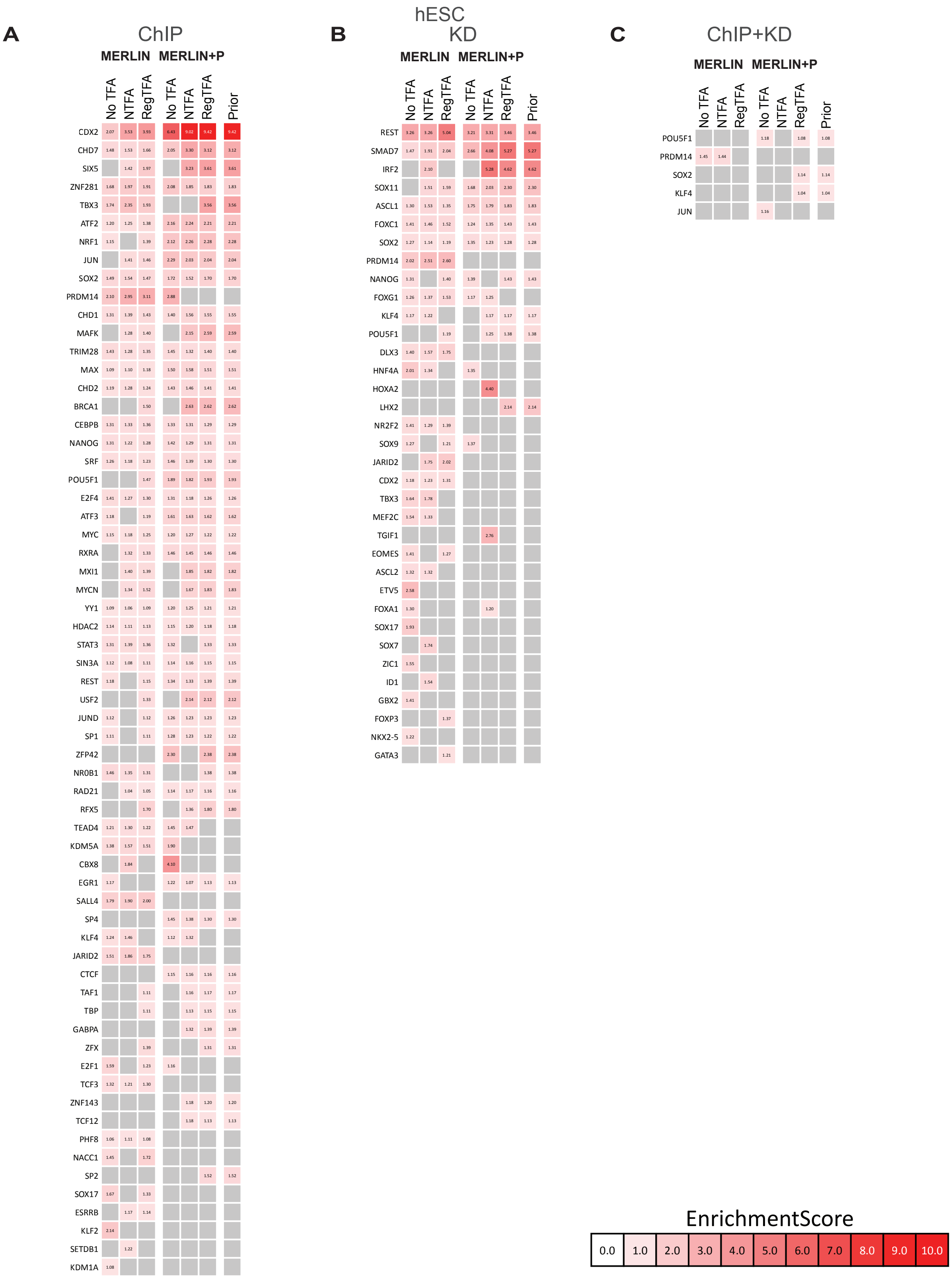
A
ChIP
MERLIN
MERLIN+P
No TFA
NTFA
RegTFA
No TFA
NTFA
RegTFA
Prior
CDX2
CHD7
SIX5
ZNF281
TBX3
ATF2
NRF1
JUN
SOX2
PRDM14
CHD1
MAFK
TRIM28
MAX
CHD2
BRCA1
CEBPB
NANOG
SRF
POU5F1
E2F4
ATF3
MYC
RXRA
MXI1
MYCN
YY1
HDAC2
STAT3
SIN3A
REST
USF2
JUND
SP1
ZFP42
NR0B1
RAD21
RFX5
TEAD4
KDM5A
CBX8
EGR1
SALL4
SP4
KLF4
JARID2
CTCF
TAF1
TBP
GABPA
ZFX
E2F1
TCF3
ZNF143
TCF12
PHF8
NACC1
SP2
SOX17
ESRRB
KLF2
SETDB1
KDM1A
B
hESC
KD
MERLIN
MERLIN+P
No TFA
NTFA
RegTFA
No TFA
NTFA
RegTFA
Prior
REST
SMAD7
IRF2
SOX11
ASCL1
FOXC1
SOX2
PRDM14
NANOG
FOXG1
KLF4
POU5F1
DLX3
HNF4A
HOXA2
LHX2
NR2F2
SOX9
JARID2
CDX2
TBX3
MEF2C
TGIF1
EOMES
ASCL2
ETV5
FOXA1
SOX17
SOX7
ZIC1
ID1
GBX2
FOXP3
NKX2-5
GATA3
C
ChIP+KD
MERLIN
MERLIN+P
No TFA
NTFA
RegTFA
No TFA
NTFA
RegTFA
Prior
POU5F1
PRDM14
SOX2
KLF4
JUN
EnrichmentScore
0.0
1.0
2.0
3.0
4.0
5.0
6.0
7.0
8.0
9.0
10.0

**Figure S4.** List of all predictable TFs when recovering **A**) ChIP, **B**) KD, and **C**) the intersection of the ChIP and KD. Each column correspond to one of inferred networks (MERLIN and MERLIN+P) using expression and TFA (at different regularization levels) of regulators. 'RegTFA' represents the best performing regularized NCA result after testing $\lambda = 0.005, 0.010, 0.020, 0.100$. The higher the enrichment score, the more significant the overlap is between the targets of that TF in that particular inferred network and the corresponding gold standard network.

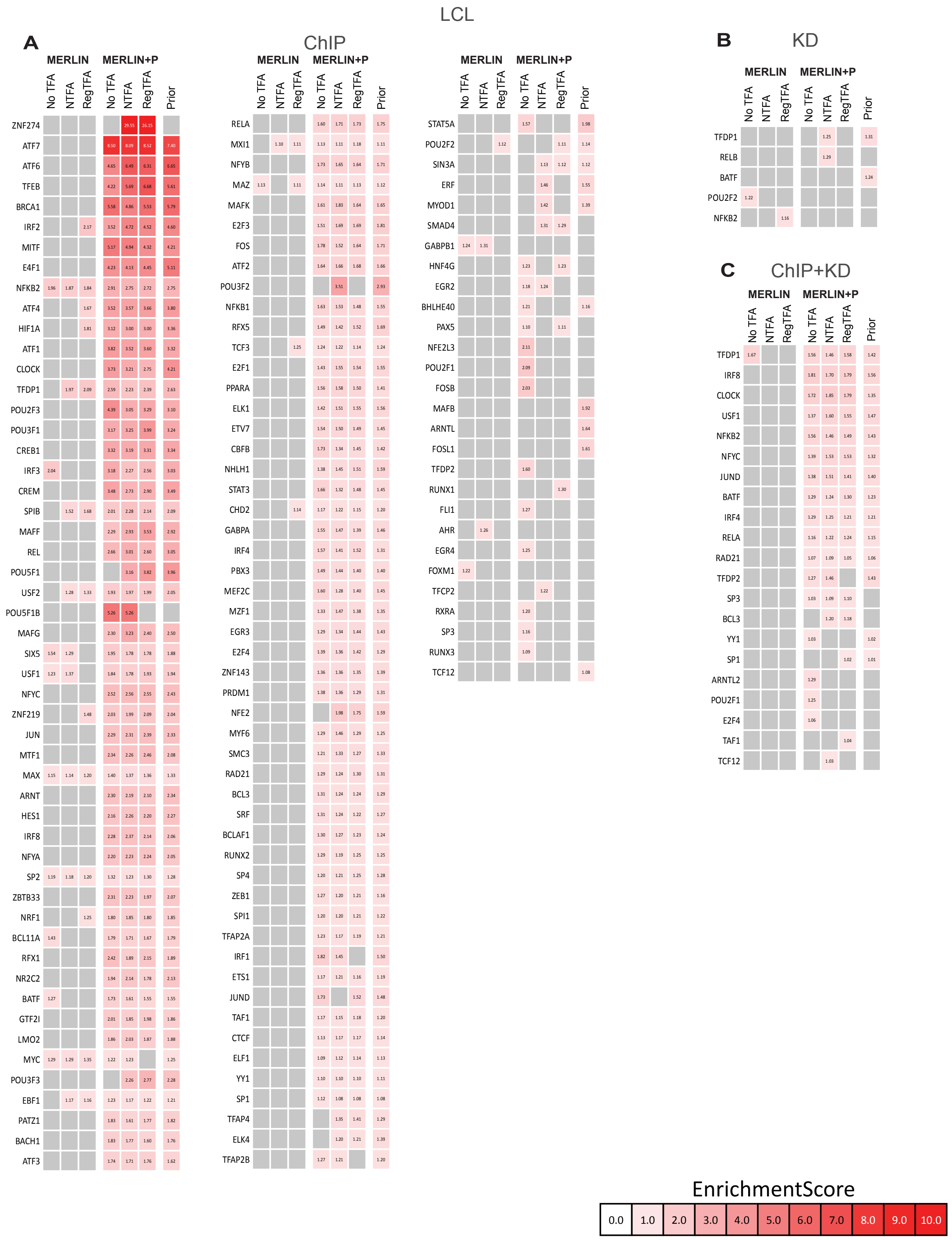
LCL
A
ChIP
MERLIN
MERLIN+P
No TFA
NTFA
RegTFA
Prior
ZNF274 ATF7 ATF6 TFEB BRCA1 IRF2 MITF E4F1 NFKB2 ATF4 HIF1A ATF1 CLOCK TFDP1 POU2F3 POU3F1 CREB1 IRF3 CREM SPIB MAFF REL POU5F1 USF2 POU5F1B MAFG SIX5 USF1 NFYC ZNF219 JUN MTF1 MAX ARNT HES1 IRF8 NFYA SP2 ZBTB33 NRF1 BCL11A RFX1 NR2C2 BATF GTF2I LMO2 MYC POU3F3 EBF1 PATZ1 BACH1 ATF3
RELA MXI1 NFYB MAZ MAFK E2F3 FOS ATF2 POU3F2 NFKB1 RFX5 TCF3 E2F1 PPARA ELK1 ETV7 CBFB NHLH1 STAT3 CHD2 GABPA IRF4 PBX3 MEF2C MZF1 EGR3 E2F4 ZNF143 PRDM1 NFE2 MYF6 SMC3 RAD21 BCL3 SRF BCLAF1 RUNX2 SP4 ZEB1 SPI1 TFAP2A IRF1 ETS1 JUND TAF1 CTCF ELF1 YY1 SP1 TFAP4 ELK4 TFAP2B
STAT5A POU2F2 SIN3A ERF MYOD1 SMAD4 GABPB1 HNF4G EGR2 BHLHE40 PAX5 NFE2L3 POU2F1 FOSB MAFB ARNTL FOSL1 TFDP2 RUNX1 FLI1 AHR EGR4 FOXM1 TFCP2 RXRA SP3 RUNX3 TCF12
B
KD
TFDP1 RELB BATF POU2F2 NFKB2
C
ChIP+KD
TFDP1 IRF8 CLOCK USF1 NFKB2 NFYC JUND BATF IRF4 RELA RAD21 TFDP2 SP3 BCL3 YY1 SP1 ARNTL2 POU2F1 E2F4 TAF1 TCF12
EnrichmentScore
0.0 1.0 2.0 3.0 4.0 5.0 6.0 7.0 8.0 9.0 10.0

**Figure S5.** List of all predictable TFs when recovering **A**) ChIP, **B**) KD, and **C**) the intersection of the ChIP and KD. Each column correspond to one of inferred networks (MERLIN and MERLIN+P) using expression and TFA (at different regularization levels) of regulators. 'RegTFA' represents the best performing regularized NCA result after testing $\lambda = 0.005, 0.010, 0.020, 0.100$. The higher the enrichment score, the more significant the overlap is between the targets of that TF in that particular inferred network and the corresponding gold standard network.

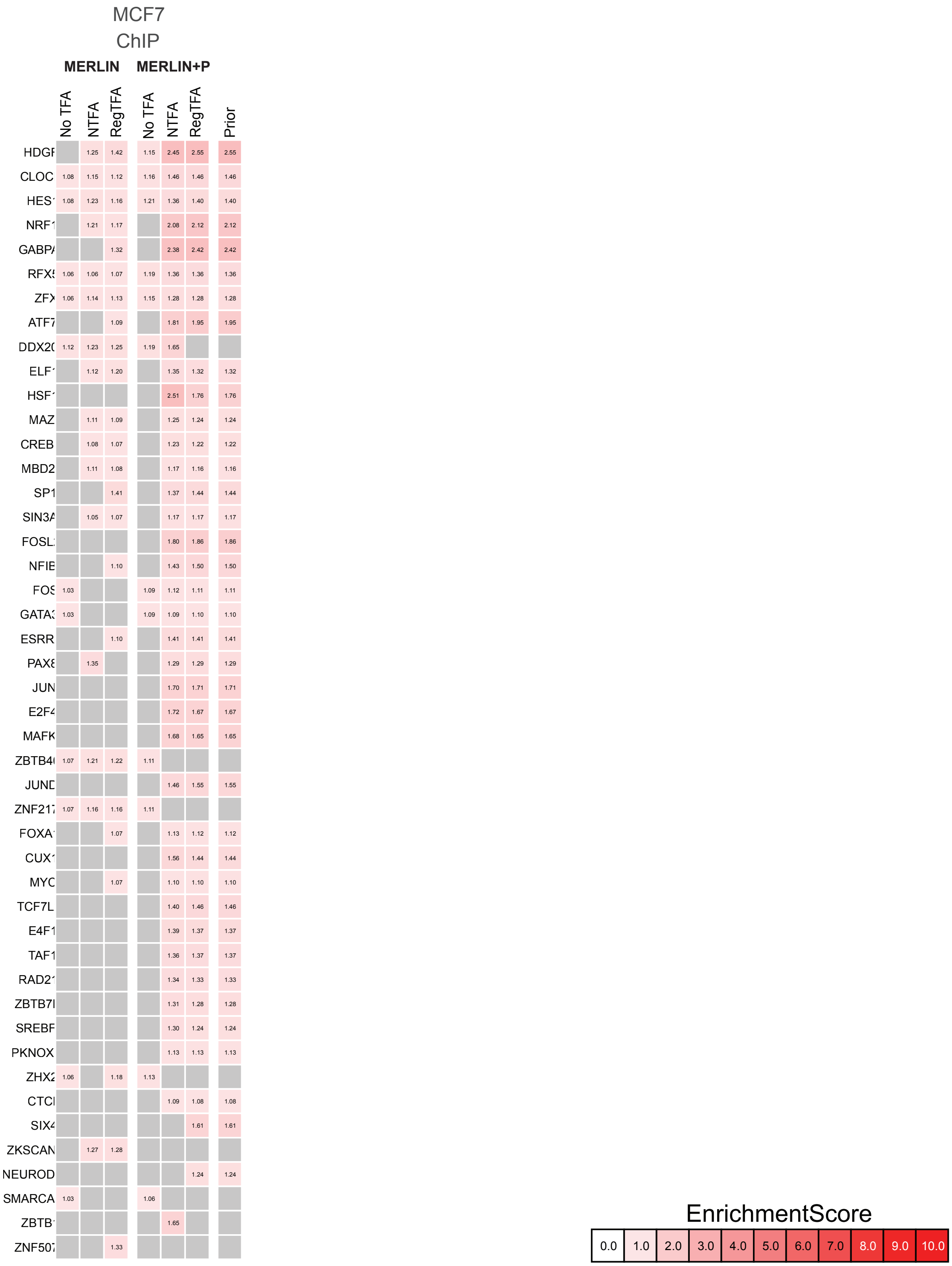
MCF7
ChIP
MERLIN
MERLIN+P
No TFA
NTFA
RegTFA
No TFA
NTFA
RegTFA
Prior
HDGF
CLOC
HES1
NRF1
GABPA
RFX5
ZFX
ATF7
DDX20
ELF1
HSF1
MAZ
CREB
MBD2
SP1
SIN3A
FOSL
NFIE
FOS
GATA3
ESRR
PAX8
JUN
E2F4
MAFK
ZBTB40
JUND
ZNF217
FOXA1
CUX1
MYC
TCF7L
E4F1
TAF1
RAD21
ZBTB7
SREBF
PKNOX
ZHX2
CTCF
SIX4
ZKSCAN
NEUROD
SMARCA
ZBTB1
ZNF507
EnrichmentScore
0.0
1.0
2.0
3.0
4.0
5.0
6.0
7.0
8.0
9.0
10.0

**Figure S6.** List of all predictable TFs when recovering ChIP. Each column correspond to one of inferred networks (MERLIN and MERLIN+P) using expression and TFA (at different regularization levels) of regulators. 'RegTFA' represents the best performing regularized NCA result after testing $\lambda = 0.005, 0.010, 0.020, 0.100$. The higher the enrichment score, the more significant the overlap is between the targets of that TF in that particular inferred network and the corresponding gold standard network.

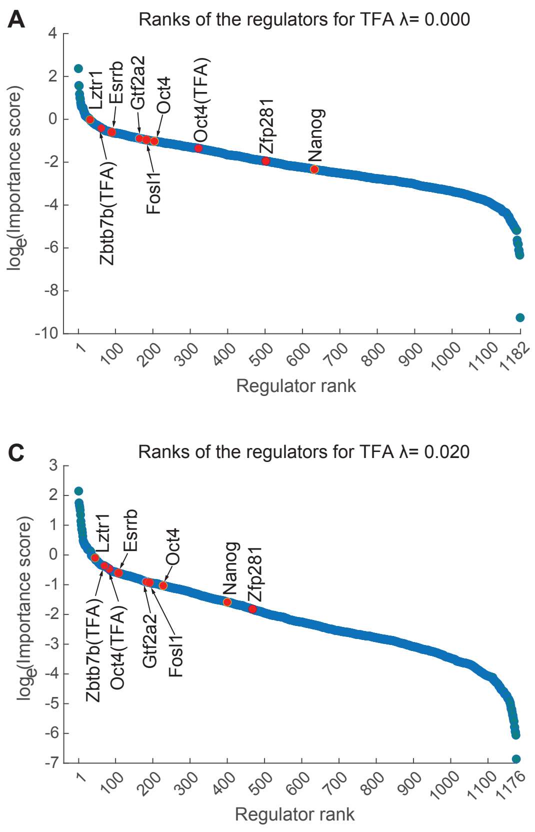
A
Ranks of the regulators for TFA λ= 0.000
loge(Importance score)
Regulator rank
Lztr1
Esrrb
Gtf2a2
Oct4
Oct4(TFA)
Zfp281
Nanog
Zbtb7b(TFA)
Fosl1
B
Ranks of the regulators for TFA λ= 0.005
Lztr1
Esrrb
Fosl1
Oct4
Zfp281
Nanog
Zbtb7b(TFA)
Gtf2a2
Oct4(TFA)
C
Ranks of the regulators for TFA λ= 0.020
Lztr1
Esrrb
Oct4
Nanog
Zfp281
Zbtb7b(TFA)
Oct4(TFA)
Gtf2a2
Fosl1
D
Ranks of the regulators for TFA λ= 0.100
Lztr1
Gtf2a2
Esrrb
Zfp281
Nanog
Oct4(TFA)
Zbtb7b(TFA)
Fosl1
Oct4

**Figure S7.** Ranked regulators in the MERLIN+P+TFA inferred network from the mESC RNA-seq dataset. Shown are the other TFA rankings ($\lambda = 0.000, 0.005, 0.020, 0.100$) besides the regularized TFA ($\lambda = 0.01$) used for prioritization of regulators for experimental validation. Four known ESC regulators (Esrrb, Nanog, Oct4, Zfp281) and 4 novel (Fosl1, Gtf2a2, Zbtb7b, Lztr1) TFs are highlighted in red cicles. "Oct4" is highlighted twice since it in the inferred GRN using gene expression (Oct4) and its estimated TFA (Oct4 (TFA)).

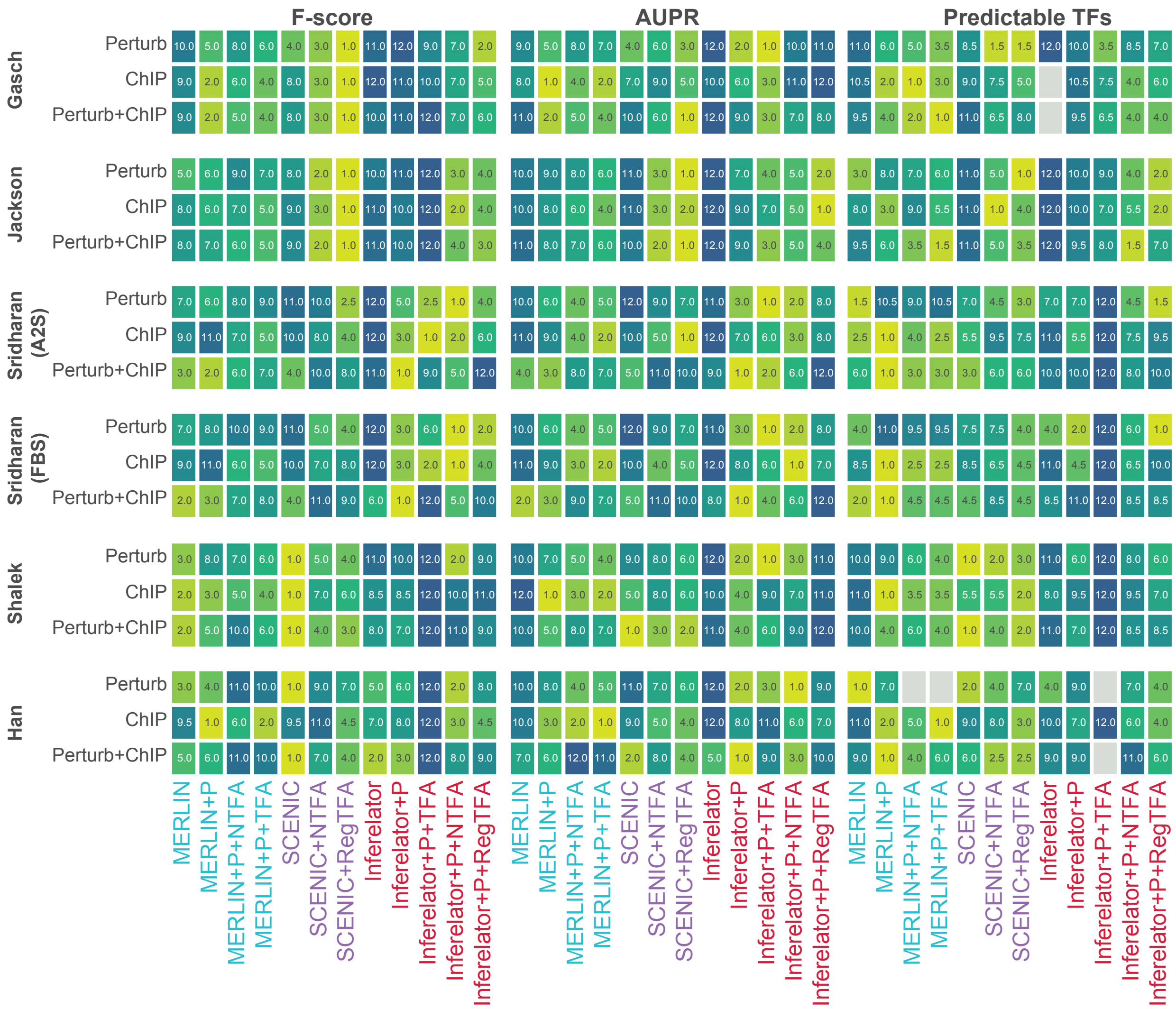

F-score
AUPR
Predictable TFs
Gasch
Jackson
Sridharan (A2S)
Sridharan (FBS)
Shalek
Han
Perturb
ChIP
Perturb+ChIP
MERLIN
MERLIN+P
MERLIN+P+NTFA
MERLIN+P+TFA
SCENIC
SCENIC+NTFA
SCENIC+RegTFA
Inferelator
Inferelator+P
Inferelator+P+TFA
Inferelator+P+NTFA
Inferelator+P+RegTFA

**Figure S8.** Algorithm performance rankings across individual scRNAseq datasets based on three gold standards: ChIP, Perturb, and their intersection (Perturb+ChIP). Performance was evaluated using F-score, AUPR, and predictable TF metrics.

**Table S1.** Abbreviations used for algorithm configurations.

| Configuration | Abbreviation | Description |
|---|---|---|
| Expression alone | No TFA | This describes an algorithm that was run without prior and/or any type of TFA. |
| MERLIN with Prior | MERLIN+P | This describes MERLIN that was run with a prior. |
| MERLIN with Prior and NCA | MERLIN+P+NTFA | This describes MERLIN that was run using a prior and NCA TFA. |
| MERLIN with Prior and Regularized TFA | MERLIN+P+TFA | This describes MERLIN that was run using a prior and regularized TFA. |
| NCA TFA | NTFA | This describes an algorithm that was run using NCA TFA. |
| Regularized TFA | RegTFA | This describes an algorithm that was run using regularized TFA. |
| NetRex inbuilt TFA | NetRex+TFA | This describes NetRex that was run using its inbuilt TFA estimation. |
| Inferelator inbuilt TFA | Inferelator+P+TFA | This describes Inferelator that was run using its inbuilt TFA estimation. |
| mLASSO inbuilt TFA | mLASSO+TFA | This describes mLASSO that was run using its inbuilt TFA estimation. |

**Table S2.** For each of the 205 regulators, we searched the relevant literature to see if there were studies reporting its involvement in pluripotency.

| Count | Regulator | Count | Regulator | Count | Regulator | Count | Regulator | Count | Regulator |
|---|---|---|---|---|---|---|---|---|---|
| 1 | Hoxa6 | 45 | Kdm5d | 89 | Isl2 | 133 | Tcfap2a | 177 | Pax5 |
| 2 | Hoxa10 | 46 | Lin54 | 90 | Jun | 134 | Tfcp2l1 | 178 | Pbx3 |
| 3 | Hoxb8 | 47 | Lztr1 | 91 | Klf4 | 135 | Tgif2 | 179 | Pim2 |
| 4 | Hoxc13 | 48 | Mlxip | 92 | Klf5 | 136 | Trp53 | 180 | Pknox2 |
| 5 | Hoxd8 | 49 | Mypop | 93 | Klf6 | 137 | Trp63 | 181 | Pou2f2 |
| 6 | Hoxd9 | 50 | Rhox6 | 94 | Max | 138 | E2f4 | 182 | Prkcq |
| 7 | Aes | 51 | Scml2 | 95 | Mbd1 | 139 | Hprt | 183 | Rfx2 |
| 8 | Alx4 | 52 | Scrt1 | 96 | Mbd2 | 140 | Ahr | 184 | Rora |
| 9 | Ankrd22 | 53 | Scrt2 | 97 | Mecom | 141 | Arnt | 185 | Rxrg |
| 10 | Arx | 54 | Six4 | 98 | Meg3 | 142 | Atf4 | 186 | Smad9 |
| 11 | Asz1 | 55 | Sp3 | 99 | Meis1 | 143 | Atf6 | 187 | Smarcc1 |
| 12 | Atf1 | 56 | Tbx20 | 100 | Mitf | 144 | Atoh1 | 188 | Sohlh2 |
| 13 | Bach1 | 57 | Tcfl5 | 101 | Mnx1 | 145 | Cdx1 | 189 | Sp1 |
| 14 | Batf3 | 58 | Whsc2 | 102 | Mtf1 | 146 | Cebpb | 190 | Spdef |
| 15 | Bhlha15 | 59 | Xbp1 | 103 | Myc | 147 | Clock | 191 | Srebf2 |
| 16 | Bhlhe40 | 60 | Zbtb7a | 104 | Myod1 | 148 | Ctcfl | 192 | Srf |
| 17 | Ccrn4l | 61 | Zbtb7b | 105 | Nfe2l2 | 149 | E2f1 | 193 | Stat3 |
| 18 | Churc1 | 62 | Zfp110 | 106 | Nfil3 | 150 | E2f2 | 194 | Tbp |
| 19 | Col5a2 | 63 | Zfp784 | 107 | Nfya | 151 | E4f1 | 195 | Tead3 |
| 20 | Creb3 | 64 | Hist1h2bb | 108 | Nhlh1 | 152 | Egr1 | 196 | Tfdp1 |
| 21 | Creb3l2 | 65 | Esrrb | 109 | Nr0b1 | 153 | En1 | 197 | Usf2 |
| 22 | Creb3l3 | 66 | Hes1 | 110 | Nr1i3 | 154 | Esr1 | 198 | Wt1 |
| 23 | Dmrta2 | 67 | Zic1 | 111 | Nr2e1 | 155 | Ets1 | 199 | Ybx1 |
| 24 | Dtx3l | 68 | Zic2 | 112 | Nr4a2 | 156 | Ets1 | 200 | Zfp128 |
| 25 | Dux | 69 | Zic3 | 113 | Nr5a2 | 157 | Etv6 | 201 | Zfp160 |
| 26 | E2f6 | 70 | Zscan4c | 114 | Nr6a1 | 158 | Foxd1 | 202 | Zfp161 |
| 27 | Ehf | 71 | Yy1 | 115 | Olig1 | 159 | Foxo4 | 203 | Zfp300 |
| 28 | Elf1 | 72 | Zeb1 | 116 | Olig2 | 160 | Foxp1 | 204 | Zfp740 |
| 29 | Enpp3 | 73 | Zfp281 | 117 | Pax2 | 161 | Gli2 | 205 | AC168977.1 |
| 30 | Fosl1 | 74 | Zfp42 | 118 | Pgr | 162 | Glis3 | | |
| 31 | Foxc1 | 75 | Zfx | 119 | Phox2b | 163 | Hlf | | |
| 32 | Foxc2 | 76 | Cd44 | 120 | Pou2f1 | 164 | Hoxa4 | | |
| 33 | Foxj3 | 77 | Ctcf | 121 | Pou3f1 | 165 | Hsf1 | | |
| 34 | Foxk2 | 78 | Eomes | 122 | Pou5f1 | 166 | Ikzf4 | | |
| 35 | Gcm2 | 79 | Fgf5 | 123 | Prrx1 | 167 | Jdp2 | | |
| 36 | Gmeb1 | 80 | Fhl2 | 124 | Rarg | 168 | Klf13 | | |
| 37 | Gmeb2 | 81 | Foxd3 | 125 | Rest | 169 | Lhx1 | | |
| 38 | Gtf2a2 | 82 | Gata2 | 126 | Smad4 | 170 | Mafk | | |
| 39 | Hes7 | 83 | Gbx2 | 127 | Snai2 | 171 | Mapk13 | | |
| 40 | Hey1 | 84 | Gli1 | 128 | Sox2 | 172 | Mecp2 | | |
| 41 | Hist1h1a | 85 | Id1 | 129 | T | 173 | Mlx | | |
| 42 | Hist1h1b | 86 | Igfbp3 | 130 | Tbx3 | 174 | Nkx2-3 | | |
| 43 | Hmbox1 | 87 | Ilk | 131 | Tcf15 | 175 | Npas3 | | |
| 44 | Hsfy2 | 88 | Irx2 | 132 | Tcf3 | 176 | Nr2c2 | | |

#### List of Files

**File S1.** Yeast all regularization settings

The file `sfile1_yeast_all_tfa_AUPR.xlsx` lists all regularization values from yeast experiments for each algorithm for each gold standard as determined by AUPR.

**File S2.** Yeast best regularization settings

The file `sfile2_yeast_best_tfa_AUPR.xlsx` lists best regularization values from yeast experiments for each algorithm for each gold standard as determined by AUPR.

**File S3.** Mammal all regularization settings

The file `sfile3_mammal_all_tfa_AUPR.xlsx` lists all regularization values from mammal experiments for each algorithm for each gold standard as determined by AUPR.

**File S4.** Mammal best regularization settings

The file `sfile5_mammal_best_tfa_AUPR.xlsx` lists best regularization values from mammal experiments for each algorithm for each gold standard as determined by AUPR.

**File S5.** Module heatmaps

The file `sfile5_merlinp_rnaseq_tfa0010_hmap_out_0_8_0_3.zip` contains module-specific heatmaps for 54 modules with enriched regulators. The folder also contains a GO enrichment and regulator enrichment bird's-eye view heatmap which presents the enriched GO terms and enriched regulators for each module.

**File S6.** KS Test P-Values

The file `sfile6_KS_test_p_vals.xlsx` contains KS test result *p*-values, histograms, and CDF plots from mESC siRNA knockdown experiments.

**File S7.** High confidence target details

The file `sfile7_high_conf_targets.xlsx` summarizes targets and gold standard support of known and novel TFs examined in validation experiments.

**File S8.** Network edge confidence

The file `sfile8_KD_edges_TFA_support.xlsx` contains edge confidence in the inferred network for targets for known and novel TFs examined during laboratory validation.

**File S9.** scRNAseq all regularization settings

The file `sfile9_scrnaseq_all_tfa_AUPR.xlsx` lists all regularization values from scRNAseq experiments for each algorithm for each gold standard as determined by AUPR.

**File S10.** scRNAseq best regularization settings

The file `sfile10_scrnaseq_best_tfa_AUPR.xlsx` lists best regularization values from scRNAseq experiments for each algorithm for each gold standard as determined by AUPR.

**File S11.** mESC ChIP gold standard

The file `sfile11_mESC_chip_rnaseq_gold.txt` lists ChIP gold standard used for mESC network comparisons.

**File S12.** mESC KD gold standard

The file `sfile12_mESC_bothko_rnaseq_gold.txt` contains KD gold standard used for mESC network comparisons.

**File S13.** MERLIN-P RNA-seq TFA 0.010 results

The file `sfile13_both.merlinp.rnaseq.tfa0.010.txt` contains the learned network inferred under TFA=0.010.

**File S14.** List of siRNA target sequences.

The file `sfile14_siRNA_sequences.xlsx` contains the details of siRNA knockdown genes.

**File S15.** Primer details.

The file `sfile15_primers.xlsx` contains primer details.
